# Excessive activation of the RAS/MAPK pathway triggers adult-onset motor axonal degeneration

**DOI:** 10.1101/2025.11.11.687669

**Authors:** Marije Been, Erwan Lambert, Ana Serna, Anouk C.G. Balvert, Simon Heisinger, Merli Kreshpani, Catherine te Dorsthorst-Maas, Paulina Paškevičiūtė, Henar Rodríguez Arias, Danique Beijer, Galuh Astuti, Kevin Kenna, Boyd van Reijmersdal, Sharon Gloudemans, Pascal van Lith, Céline Sijlmans, Patrik Verstreken, Jan Veldink, Stephan Züchner, Christian Gilissen, Annette Schenck, Erik Storkebaum

**Affiliations:** Molecular Neurobiology Laboratory, Donders Institute for Brain, Cognition and Behaviour and Faculty of Science, Radboud University, Nijmegen, Netherlands; Dr. John T. Macdonald Foundation Department of Human Genetics and John P. Hussman Institute for Human Genomics, University of Miami Miller School of Medicine, Miami, FL 33136, USA; Department of Human Genetics, Radboud Institute for Molecular Life Sciences, Radboud University Medical Center, Nijmegen, Netherlands; Department of Neurology, UMC Utrecht Brain Centre, University Medical Centre Utrecht, Utrecht, Netherlands; Department of Human Genetics, Donders Institute for Brain, Cognition and Behaviour, Radboud University Medical Center, Nijmegen, Netherlands; VIB-KU Leuven Center for Brain & Disease Research, Leuven 3000, Belgium; Department of Neurosciences, Leuven Brain Institute, KU Leuven, 3000 Leuven, Belgium

**Author notes:** These authors contributed equally (Anouk C.G. Balvert, Simon Heisinger, Merli Kreshpani, Catherine te Dorsthorst-Maas, Paulina Paškevičiūtė, Henar Rodríguez Arias). Corresponding authors: Marije Been, Erik Storkebaum.

## Abstract

Axonal and synaptic degeneration are key hallmarks of neurodegenerative diseases, but the underlying molecular mechanisms are incompletely understood. Here, we performed an unbiased forward genetic mosaic screen to identify genes required for maintenance of adult motor axons and neuromuscular junctions (NMJs) in the *Drosophila* leg. We identified 49 mutations in 30 genes, including mutations in 8 genes resulting in adult-onset progressive degeneration. We found that loss of *pebbled* (*peb*) function results in adult-onset motor axonal and NMJ degeneration, and age-dependent motor deficits. *Peb* is the *Drosophila RREB1* ortholog, a C_2_H_2_ zinc-finger transcription factor that negatively regulates transcription of RAS/MAPK pathway target genes. Loss of *peb* function resulted in excessive RAS/MAPK pathway activation, and loss of function of other negative regulators of the RAS/MAPK pathway also induced adult-onset progressive NMJ degeneration and motor deficits. Importantly, treatment of adult flies with the MEK1/2 inhibitor mirdametinib induced a dosage-dependent rescue of *peb* mutant motor neurodegenerative phenotypes. Thus, RAS/MAPK pathway overactivation results in adult-onset progressive neurodegeneration, which can be prevented by RAS/MAPK pathway inhibition.

## Introduction

Neurodegenerative disorders are typically adult-onset and progressive diseases, characterized by degeneration of specific neuronal populations. Axonal degeneration is a prominent pathological feature of many neurodegenerative diseases, and axons and presynaptic terminals are often the neuronal compartments first affected (Wilson *et al*, 2023). Neurodegenerative diseases have a high and increasing prevalence in the aging population, have devastating consequences for patients and their families, and result in high costs for society. The molecular mechanisms underlying neurodegenerative diseases are incompletely understood and, as a consequence, no or only minimally effective disease-modifying treatments are available for the vast majority of neurodegenerative diseases. Thus, there is an urgent need to gain deeper insight into the molecular mechanisms underlying neurodegeneration.

Here, we performed an unbiased, forward genetic mosaic screen to identify genes on the *Drosophila* X chromosome that are required for axonal maintenance. EMS-based forward genetic screens in *Drosophila* have been instrumental in uncovering the molecular mechanisms underlying key biological processes, including embryonic polarity (Nüsslein-volhard & Wieschaus, 1980), synaptic function (Verstreken *et al*, 2003; Stowers *et al*, 2002; Dickman & Davis, 2009), and neuronal development, function and maintenance (Yamamoto *et al*, 2014; Chihara *et al*, 2007; Pielage *et al*, 2008). A large-scale genetic screen for genes with essential functions in neuronal development, function, and maintenance identified 165 genes that when mutated resulted in defects in mechanosensory organs (bristles), wing morphogenesis, eye/head morphology, or functional electroretinogram (ERG) defects (Yamamoto *et al*, 2014). This screen provided valuable insights into molecular mechanisms underlying several neurological disorders (Goodman *et al*, 2021; Fishburn *et al*, 2024; Pan *et al*, 2024).

The screen presented here has a unique design which allowed for the identification of gene mutations resulting in adult-onset degeneration of peripheral motor and sensory axons and neuromuscular junctions (NMJs) in the *Drosophila* leg. Our screen identified 30 genes that are involved in various molecular processes. 29 (97%) of the identified genes have human orthologs, of which 19 (66%) have orthologs that have previously been associated with neurodegenerative and neurodevelopmental disorders, confirming the disease relevance of our screen.

One of the identified genes is *pebbled* (*peb*), the *Drosophila* ortholog of human *RAS-responsive element binding protein 1* (*RREB1*). *Peb* and *RREB1* encode C_2_H_2_ zinc-finger transcription factors, and RREB1 negatively regulates transcription of RAS-MAPK pathway target genes through binding to RAS-response elements (RREs) in their promoters (Kent *et al*, 2020). Haploinsufficiency of *RREB1* causes a Noonan-like RASopathy in humans and mice (Kent *et al*, 2020; Shatokhina *et al*, 2024; Strong *et al*, 2025). RASopathies, including Noonan syndrome, affect ∼1/1,000 individuals and are characterized by short stature, facial dysmorphism, intellectual disability and cardiac abnormalities (Zenker, 2022; Roberts *et al*, 2013). RASopathies are caused by germline mutations in genes encoding components or regulators of the RAS-MAPK pathway (Tartaglia *et al*, 2022; Rauen, 2013; Hebron *et al*, 2022), either activating mutations in RAS/MAPK pathway components, or loss-of-function mutations in negative regulators of the RAS/MAPK pathway. The latter category includes neurofibromatosis type 1 (NF1), Legius syndrome, and *LZTR1*-associated Noonan syndrome. NF1 is caused by heterozygous loss-of-function mutations in the *NF1* gene, which encodes neurofibromin, a RAS GTPase-activating protein (GAP) that negatively regulates RAS by stimulating the conversion of active RAS-GTP to inactive RAS-GDP (Rauen, 2013). Legius syndrome is caused by heterozygous inactivating mutations in *SPRED1*, a negative regulator of the RAS/MAPK pathway by inhibiting phosphorylation of Raf (Rauen, 2013; Zenker, 2022; Roberts *et al*, 2013). Heterozygous and homozygous *LZTR1* mutations can cause Noonan syndrome. LZTR1 functions as substrate receptor for cullin3 (CUL3)-RING ubiquitin ligase (CRL3) complexes (Motta *et al*, 2019). Because LZTR1 substrates include RAS and RIT1, LZTR1 negatively regulates the RAS/MAPK pathway by enabling RAS and RIT1 ubiquitination (Bigenzahn *et al*, 2018). The Noonan syndrome causative mutations impede LZTR1 substrate recognition, thus resulting in excessive RAS/MAPK pathway activation. Interstingly, beyond RASopathies, excessive activation of the RAS/MAPK pathway was suggested to be implicated in neurodegenerative diseases, including Parkinson’s disease (Reinhardt *et al*, 2013) and amyotrophic lateral sclerosis (ALS) (Ayala *et al*, 2011a; Caldi Gomes *et al*, 2024), but *in vivo* evidence for a causative link between the RAS/MAPK pathway and neurodegeneration is missing.

We discovered that loss of function of *peb*, as well as other negative regulators of the RAS/MAPK pathway such as *Neurofibromin 1* (*Nf1*), *Spred, and Lztr1,* caused adult-onset progressive degeneration of motor axons and NMJs and resulted in age-related, progressive motor deficits. These findings identify overactivation of the RAS/MAPK pathway as a novel molecular mechanism underlying adult-onset progressive neurodegeneration. Importantly, the *peb* mutant axonal degeneration phenotype, motor deficit and the downstream molecular defects in α-tubulin acetylation and autophagy could be fully rescued by the small molecule MEK1/2 inhibitor mirdametinib, suggesting that pharmacological inhibition of the RAS/MAPK pathway may be explored as a novel therapeutic approach to mitigate progressive axonal degeneration in RASopathies and neurodegenerative diseases.

## Results

### An unbiased forward genetic screen of the *Drosophila* X chromosome identifies genes required for maintenance of peripheral motor axons

To identify essential genes required for axonal maintenance, we performed an unbiased, forward genetic mosaic screen of the *Drosophila* X chromosome. Using a low concentration of the chemical mutagen <u>e</u>thyl <u>m</u>ethane<u>s</u>ulfonate (EMS) to introduce random genomic mutations on a FRT19A chromosome, we established 8,892 stocks, of which 1,416 displayed hemizygous male lethality (Figure 1A, Figure S1A-D). This corresponds to a saturation of approximately 88% of the X chromosome (see Methods). To generate GFP-labeled homozygous mutant motor and sensory neuron clones in otherwise heterozygous flies, we applied mosaic analysis with a repressible cell marker (MARCM)(Lee & Luo, 1999), using *asense-FLP* (Sreedharan *et al*, 2015; Neukomm *et al*, 2014), and *OK371-GAL4* to express membrane-tethered GFP (UAS-mCD8::GFP) in glutamatergic neurons, which include motor and sensory neurons in the *Drosophila* leg (Figure 1B, Figure S1E). We screened the legs of at least two 21-day-old flies per mutant for any motor neuron morphology defects using fluorescence microscopy (Figure S1F). This approach identified motor axonal and NMJ phenotypes in 130 mutant lines, including 58 lines in which no GFP-positive motor or sensory neurons were visible at day 1 and 21. These 58 lines presumably contain cell lethal mutations and were kept for mapping purposes (Figure S2, Table S1). In the remaining 72 mutants, both motor axonal and NMJ phenotypes were present (Figure 1C for representative examples).

**Figure 1:**
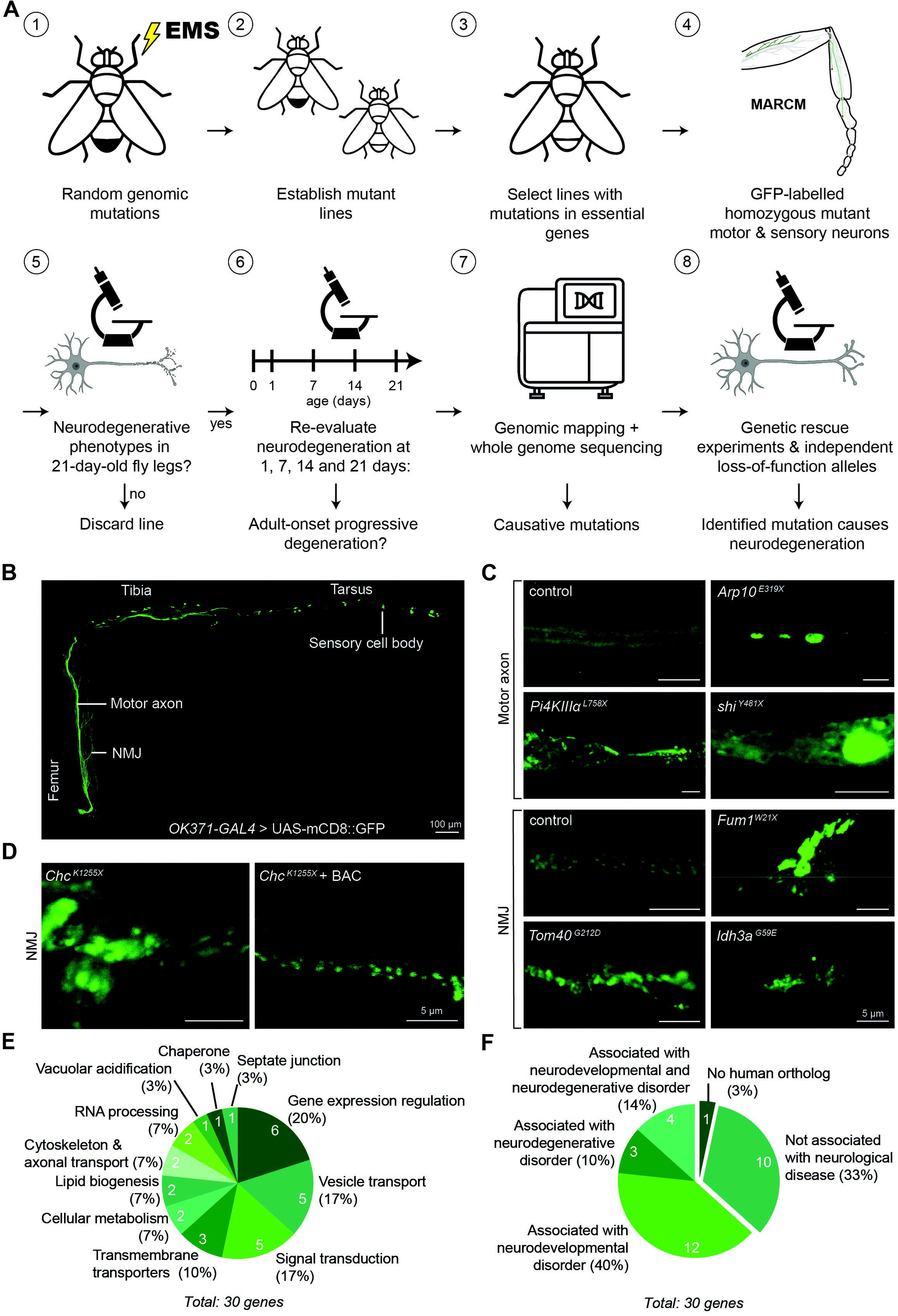
An unbiased forward genetic screen of the *Drosophila* X chromosome identifies genes involved in axonal maintenance. **(A)** Flow chart of the genetic screening approach. See text for details. **(B)** Visualization of glutamatergic neurons in the fly leg, using *OK371-GAL4* to drive expression of membrane-bound GFP (mCD8::GFP). In the leg, glutamatergic neurons include motor neurons innervating leg muscles and sensory neurons projecting to the central nervous system. Scale bar: 100 μm**. (C)** Representative examples of motor axonal and NMJ degenerative phenotypes as compared to control (FRT19A). **(D)** Phenotypic rescue of *Chc^K1255X^* NMJ phenotype by re-introduction of wild type *Chc* in a BAC-transgene. Scale bars in C,D: 5 μm. **(E,F)** Pie charts showing (E) the molecular function and (F) the disease-association of the genes identified in the screen.

Duplication and complementation mapping were performed, along with whole genome sequencing to identify the causative mutations (Figure S1G, Figure S2, Table S1). To confirm that the identified gene mutation is responsible for the neuronal morphology defect, we evaluated whether a BAC transgene containing the wild-type gene rescued the degenerative phenotype (Figure 1D for a representative example, Table S1). By using this approach, we identified 49 causative mutations in 30 genes (Table 1, Table S1). These genes are involved in diverse molecular processes (Figure 1E). Many of these pathways, such as gene expression regulation, vesicle transport and cellular metabolism, have already been associated with neurodegenerative processes (Wilson *et al*, 2023). In contrast, some of the identified pathways have not previously been related to neuronal phenotypes. For example, the link between *Drosophila* smooth septate junctions and neuronal morphology remains to be characterized.

**Table 1:**
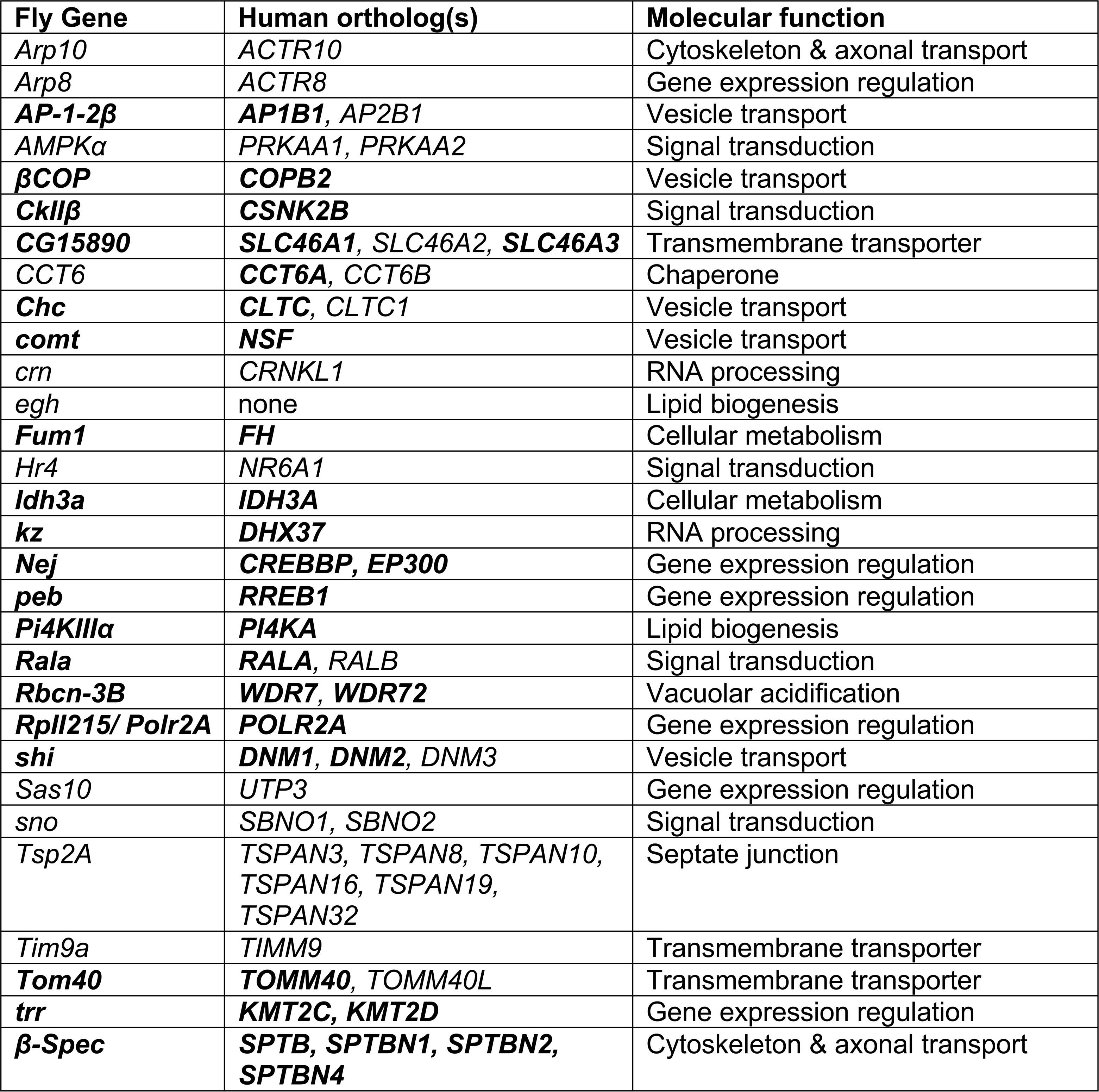
List of fly genes identified in the screen, their human orthologs and the molecular process they are involved in. See also Table S2. Human genes and the corresponding fly genes that are associated with disease are shown in bold.

29 (97%) of the identified genes have at least one human ortholog, corresponding to a total of 51 orthologous human genes. 19 (66%) of the *Drosophila* genes have human orthologs that have been associated with neurological disease (Figure 1F, Table 1, Table S2). By screening aged flies, we aimed to enrich for genes involved in neurodegeneration rather than genes involved in neuronal development. Surprisingly, only 3 (10%) of the identified *Drosophila* genes have orthologs that have exclusively been associated with neurodegenerative disease, while another 4 (14%) genes have orthologs that have been associated with both neurodevelopmental and neurodegenerative disorders. Furthermore, 12 (40%) genes have orthologs associated with neurodevelopmental disorders (Figure 1F, Table S2). Interestingly, even though our screen was designed to identify mutations that cause neurodegeneration through a recessive loss-of-function mechanism, 63% (22 out of 35) of the neurological disorders associated with the human orthologs display an autosomal dominant inheritance pattern, compared to 37% (13 out of 35) with a recessive inheritance pattern. This implies that the human orthologs of the 10 *Drosophila* genes that have hitherto not been associated with neurological disease may represent candidate genes for both recessive and dominantly inherited neurological disorders. 12 (40%) of the 30 genes had not been identified in the X chromosome screen by Yamamoto et al (Yamamoto *et al*, 2014), indicating that our unique screening strategy for adult-onset progressive degeneration was successful in identifying novel axonal degeneration genes, while confirming others.

### Phenotypic characterization of the identified axonal degeneration mutants

We next phenotypically characterized the identified mutant lines in more detail. Lethal stage determination of the 46 non-cell lethal lines revealed that hemizygous males display embryonic lethality in 23 (50%) lines, larval lethality in 21 (46%) lines, and pupal lethality in the remaining 2 (4%) lines (Table S3). Remarkably, for only 5 (18%) of 28 axonal degeneration genes evaluated, transgenic RNAi-mediated knockdown in motor neurons (*OK371-GAL4*) induced a neurodegenerative phenotype similar to the mutant (Table S3). For 6 (21%) of the identified genes, transgenic RNAi knockdown induced a very mild phenotype, whereas for the remaining 17 (61%) genes RNAi knockdown did not induce axonal or NMJ morphological phenotypes. This is likely attributable to the fact that RNAi-mediated knockdown is not complete and therefore induces partial loss-of-function, whereas homozygous damaging mutations may induce full loss-of-function. These findings emphasize the value of EMS screens as a complementary approach to transgenic RNAi screens.

To discriminate between developmental and adult-onset phenotypes, we repeated the MARCM analysis for the identified non-cell lethal mutant lines (46 mutants) and examined motor axonal and NMJ morphology at 1, 7, 14 and 21 days of age (Figure 2A, Table S3). Mutants were categorized as either 1) developmental non-progressive when the phenotype was present on day 1 and did not become more pronounced with ageing, 2) developmental progressive when a phenotype was present on day 1 and progressed with ageing, and 3) adult-onset progressive when phenotype onset was after day 1 (Figure 2A for representative examples). While the majority of mutants displayed a developmental phenotype, mutations in 8 (29%) of the 28 non-cell lethal genes resulted in an adult-onset progressive phenotype (Figure 2B, Table S3). Remarkably, mutations in the majority of the 15 non-cell lethal genes of which the human orthologs are associated with neurodevelopmental disorders resulted in progressive neurodegeneration (Figure 2C). Indeed, 9 (60%) of the 15 genes showed a developmental progressive phenotype, while only 4 (27%) showed a developmental non-progressive phenotype. Mutations in 2 genes, Ras-like protein A (*Rala*) and pebbled (*peb*), which are known to regulate neurodevelopment in other contexts (Kent *et al*, 2020; Oliva & Sierralta, 2010; Hiatt *et al*, 2018), showed an adult-onset progressive phenotype in the NMJ. These data suggest that many neurodevelopmental genes also play an important role in axonal maintenance.

**Figure 2:**
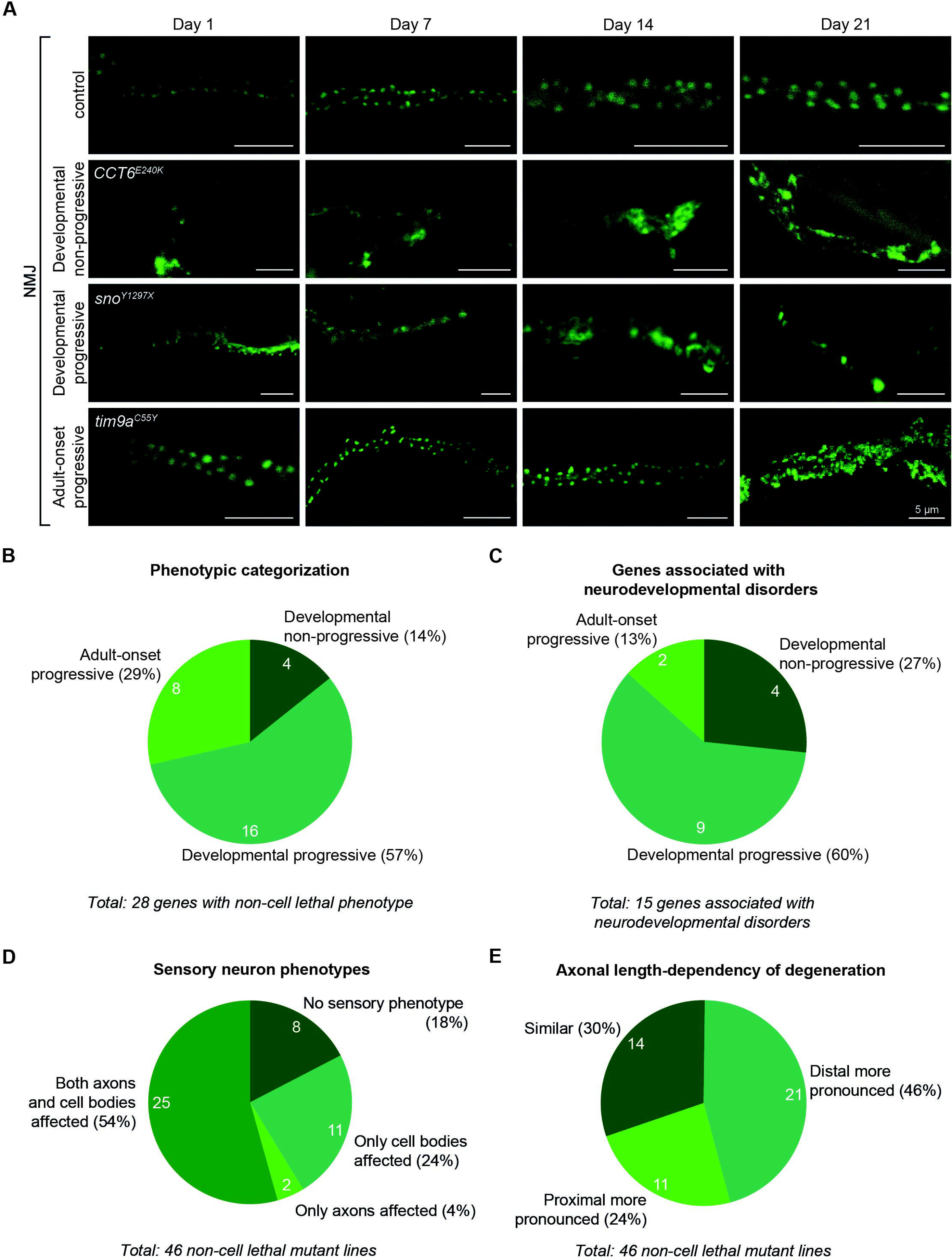
Phenotypic categorization of the identified axonal degeneration mutants. **(A)** Time course of NMJ phenotypes in the femur of FRT19A control and representative mutants. Three phenotypic categories were distinguished: (i) phenotypes present on day 1 that do not progress with ageing are classified as developmental non-progressive (e.g. *CTT6^E240K^*); (ii) developmental phenotypes that degenerate further with age are developmental progressive (e.g. *sno^Y1297X^*); (iii) mutants with a phenotype onset after day 1 are adult-onset progressive (e.g. *tim9a^C55Y^*). Scale bars: 5 μm. **(B,C)** Pie charts showing the time course phenotypic categorization of (B) all identified non-cell lethal genes and (C) mutants in genes associated with neurodevelopmental disorders. **(D)** Pie chart showing the sensory neuron phenotypes of the identified non-cell lethal mutant lines. **(E)** Pie chart showing axonal length-dependent degeneration characteristics of the identified non-cell lethal mutant lines, evaluated by comparing distal (tibia) versus proximal (femur) motor axonal and NMJ phenotypes.

We further assessed all mutants for morphological phenotypes in sensory cell bodies and axons in the *Drosophila* leg (Figure 2D, Table S3). Interestingly, 8 (17%) of the 46 non-cell lethal mutants, including mutations in *Arp8*, *CkIIβ*, *CCT6*, *Hr4*, *Idh3a*, *Polr2A, peb* and *Sas10*, did not show sensory neuron morphological defects, suggesting their selective requirement for motor neuron maintenance. The majority of the remaining mutants displayed morphological changes in both sensory axons and cell bodies, but in some mutants the sensory neuron cell body or axon were selectively affected (Figure 2D, Table S3). Finally, we compared the phenotypic severity of motor axon and NMJ degeneration in the femur and tibia, to evaluate possible axonal length-dependent degeneration (the tibia is distal to the femur). This revealed that about half of the non-cell lethal mutants (21 out of 46, 46%) displayed more pronounced phenotypes in axons and NMJs innervating distal muscles, while 30% (14 out of 46) of mutants displayed similar phenotypic severity in femur and tibia, and 24% (11 out of 46) displayed more pronounced proximal defects (Figure 2E, Table S3).

### Loss of *pebbled* function results in adult-onset progressive motor neuron degeneration

One of the mutants we identified carries a nonsense mutation in *pebbled* (*peb^Q894X^*) (Figure 3A, Figure S3A), a C_2_H_2_ zinc-finger transcription factor that is orthologous to human *RAS-responsive element binding protein 1* (*RREB1*). Homozygosity for *peb^Q894X^* resulted in adult-onset progressive degeneration of motor axons and NMJs (Figure 3B-D). We observed the accumulation of GFP-positive puncta (labeled by membrane-bound mCD8::GFP) in motor axons by day 7 whereas most NMJs were degenerated by day 21. In agreement with Farley et al (Farley *et al*, 2018), we found that loss of *peb* function does not affect sensory neuron morphology (Figure 3E). To exclude that the phenotypes are caused by a second-site mutation, we introduced an 80 kb BAC transgene containing a wild-type copy of *peb* (*peb^BAC^*), which fully rescued *peb^Q894X^* motor axonal and NMJ phenotypes (Figure 3F,G). Furthermore, we assessed motor axon and NMJ morphology for two additional *peb* loss-of-function alleles, *peb^E8^* (Strecker *et al*, 1991) and *peb^345x^*(Farley *et al*, 2018) (Figure 3B-D). Both alleles displayed a similar phenotype as the *peb^Q894X^* mutant, albeit with an earlier onset of NMJ degeneration, and were fully rescued by *peb^BAC^* (Figure 3G). This suggest that *peb^Q894X^* is a partial loss-of-function (hypomorphic) allele, whereas *peb^E8^* and *peb^345x^*were reported to be full loss-of-function (amorphic) alleles (Yip *et al*, 1997; Drysdale *et al*, 1993; Farley *et al*, 2018).

**Figure 3:**
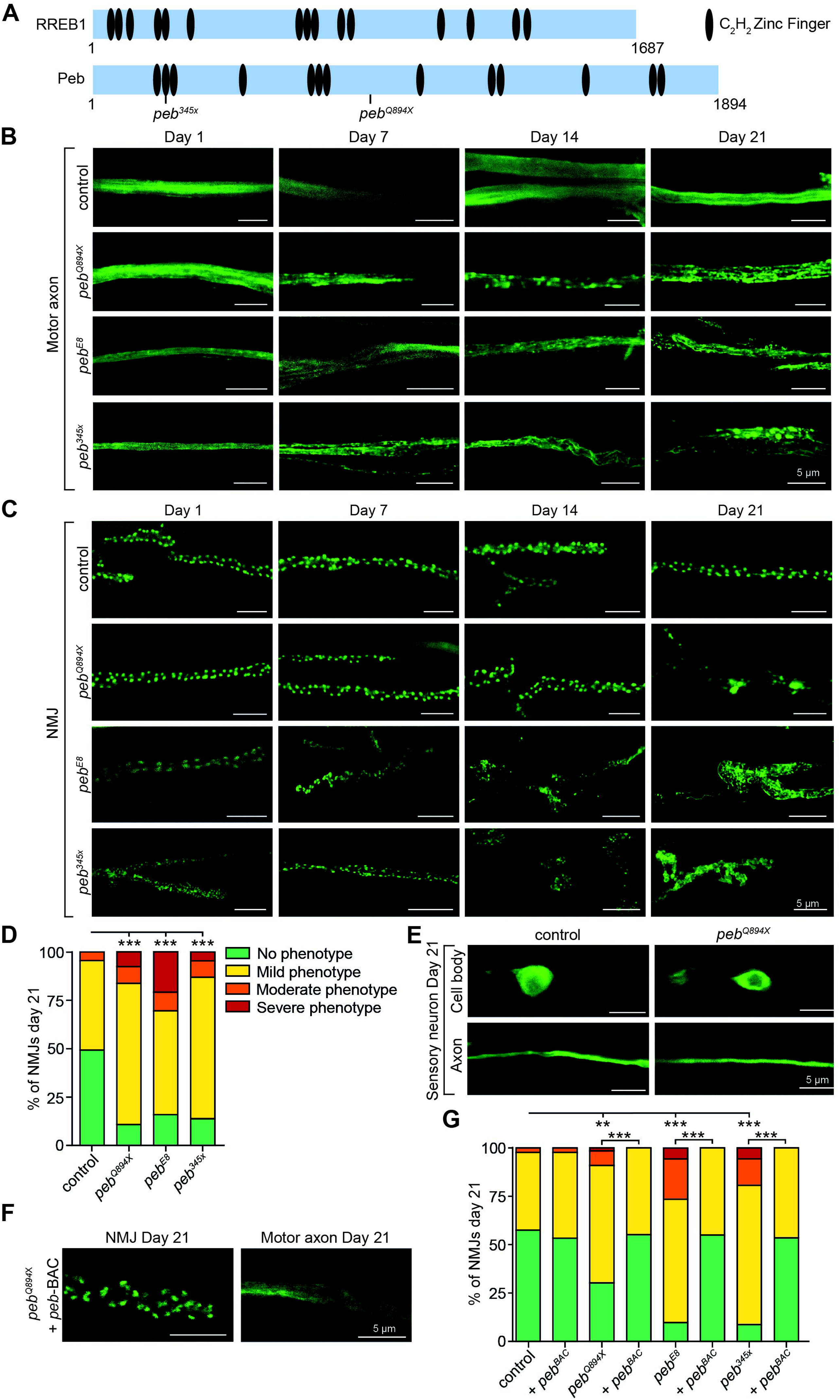
Loss of *pebbled* function results in adult-onset neurodegeneration. **(A)** Schematic representation of Peb protein and its human ortholog RREB1, showing the location of the C_2_H_2_ zinc finger domains and the *peb^345x^*and *peb^Q894X^* mutations. **(B,C)** Time course of (B) motor axon and (C) NMJ morphology in the femur of *peb^Q894X^*, *peb^E8^* and *peb^345x^* mutants versus control (FRT19A). **(D)** Semi-quantitative analysis of NMJ degeneration in the femur of three independent *peb* mutants versus control (FRT19A). NMJ phenotype was scored as no, mild, moderate or severe degeneration as illustrated in Figure 3C. **(E)** Representative images of cell bodies and axons of control (FRT19A) and *peb^Q894X^* sensory neurons at 21 days of age. **(F)** Representative images and **(G)** semi-quantitive analysis of rescue of *peb* mutant NMJ and axon morphology defects at 21 days of age by a BAC transgene containing the wild type *peb* gene (*peb*-BAC). **P<0.01, ***P<0.0001 by (D,G) Fisher’s exact test with Bonferroni correction. n= (D) 38-123 NMJs of 6-10 animals per genotype and (G) 71-107 NMJs of 6-10 animals per genotype. Data are represented as mean ± SEM. Scale bars: 5 μm.

To evaluate the effect of the *peb^Q894X^* mutation on Peb protein levels, we performed immunohistochemistry using the monoclonal anti-Peb antibody 1G9 (DSHB) and quantified the staining intensity in motor neuron cell bodies of homozygous mutant versus wild type MARCM clones in the adult ventral nerve cord (VNC). Peb staining intensity in homozygous *peb^Q894X^* clones was significantly reduced by ∼80% as compared to control (Figure 4A, representative images in Figure S3B). Because the 1G9 epitope is between amino acids 824 and 1125, this Peb antibody may still detect truncated Peb protein produced by the *peb^Q894X^* allele. To evaluate this possibility, we performed fluorescent *in situ* hybridization on VNCs of 21-day-old *peb^Q894X^* and *peb^E8^* MARCM flies. We found that *peb^Q894X^* mRNA levels in homozygous mutant motor neuron clones were not significantly reduced as compared to control, whereas *peb^E8^* mRNA levels were very low (Figure 4B, representative images in Figure S3C). Thus, it is conceivable that the milder *peb^Q894X^*neurodegenerative phenotype is due to the production of truncated Peb protein which may be partially functional.

**Figure 4:**
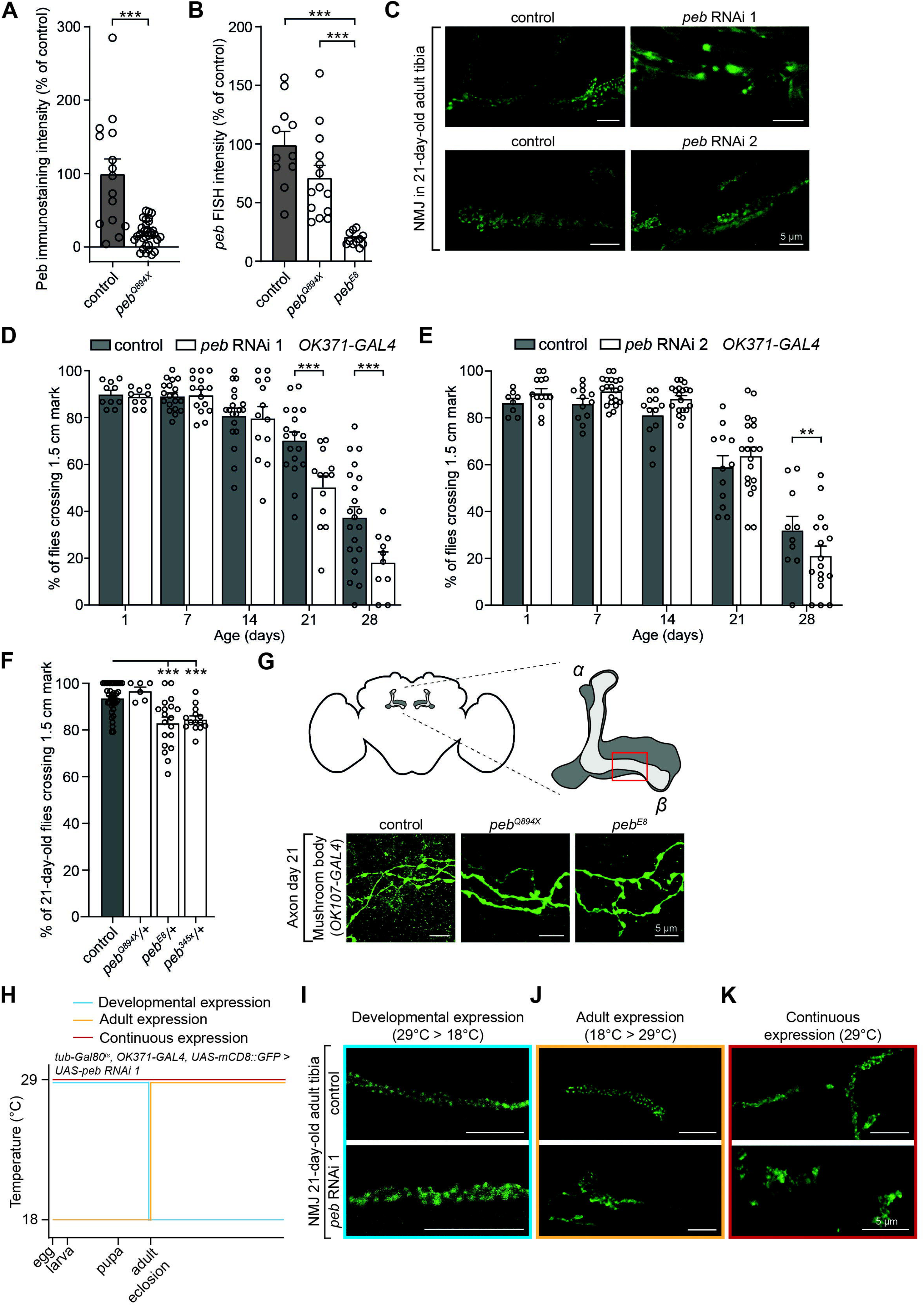
Partial loss of *peb* function induces NMJ degeneration and late-onset motor deficits. **(A)** Quantification of Peb immunostaining intensity in cell bodies of control (FRT19A) versus *peb^Q894X^* motor neuron clones in adult VNC at 21 days of age. Data are shown as percentage of control. **(B)** Quantification of signal intensity of Fluorescent *In Situ* Hybridization (FISH) for peb transcript in cell bodies of *peb* mutant motor neuron clones in the adult VNC at 21 days of age. Data are shown as percentage of control (FRT19A). **(C)** Representative images of tibial NMJs of 21-day-old flies with *peb* knockdown in motor neurons (*OK371-GAL4*). **(D-F)** Motor performance in a negative geotaxis climbing assay, shown as percentage of flies crossing a 1.5 cm mark. (D,E) Flies with *peb* knockdown in motor neurons (*OK371-GAL4*) are compared to control (KK background) at 1, 7, 14, 21, and 28 days of age using two independent RNAi lines. (F) Flies with heterozygous mutations in *peb* are compared to control (FRT19A) at 21 days of age. **(G)** Representative images of axons in control (FRT19A) and *peb* mutant MARCM clones in the β-lobe of the mushroom body (*OK107-GAL4*). Axons were imaged within the area indicated by the red square. **(H)** Schematic representation of experimental approach and **(I-K)** representative images of tibial NMJs in 21-day-old animals with *peb* knockdown in motor neurons (*OK371-GAL4*), either only during development (I), adulthood (J), or both (K). A temperature-sensitive GAL80 transgene was used to gain temporal control over *peb* RNAi transgene expression. **P<0.01, ***P<0.005 by (A) unpaired t-test with Welch’s correction, (B,F) Brown-Forsythe and Welch ANOVA with Dunnett’s T3 multiple comparisons test and (D,E) two-way ANOVA with Šidák’s multiple comparisons test. n= (A) 15-31 cells in 3 animals per genotype, (B) 11-14 animals per genotype, (D) 9-21 groups of 10 flies per genotype, (E) 8-20 groups of 10 flies per genotype, (F) 6-41 groups of 5-10 flies per genotype. Data are represented as mean ± SEM. Scale bars: 5 μm.

We next evaluated whether transgenic RNAi-mediated knockdown of *peb* would be sufficient to mimic the mutant phenotype. Expression of two independent *peb* RNAi transgenes in motor neurons (*OK371-GAL4*) resulted in NMJ degeneration in the tibia of 21-day-old animals (Figure 4C), with *peb* RNAi 1 inducing more pronounced defects. To determine whether the NMJ morphology defects observed in *peb* knockdown animals resulted in motor performance deficits, we performed a negative geotaxis climbing assay (Niehues *et al*, 2015). Both *peb* RNAi lines resulted in significant motor performance defects from 21 (*peb* RNAi 1) or 28 (*peb* RNAi 2) days-of-age onwards (Figure 4D,E). In addition, heterozygous *peb^E8/+^*and *peb^345x/+^*, but not *peb^Q894X/+^* females, displayed motor performance deficits at 21 days of age (Figure 4F). Thus, loss of *peb* function results in progressive motor axonal and NMJ degeneration, as well as motor deficits.

Because loss of a nervous system-enriched *Rreb1* transcript (*V7* isoform) in *Rreb1-nm3888^-/-^* mice was recently reported to induce progressive loss of cerebellar Purkinje cells (Griffin *et al*, 2024), whereas no phenotypes were observed in *peb* mutant sensory neurons in *Drosophila* wings and legs, we asked whether other neuronal cell types in *Drosophila* would show progressive degeneration upon loss of *peb* function. *Peb* homozygous mutant clones of both the hypomorphic *peb^Q894X^* and amorphic *peb^E8^* in mushroom body β-lobe neurons did not display axonal degeneration in 21-day-old flies (Figure 4G), despite the fact that Fly Cell Atlas, a single nucleus transcriptomic atlas of the adult fruit fly (Li *et al*, 2024), shows *peb* co-expression with known mushroom body markers *FasII*, *drk*, *Pka-R2* and *dco* (Crittenden *et al*, 1998). Together, these data suggest that motor neurons are particularly sensitive to loss of *peb* function in *Drosophila*.

Next, we wanted to confirm that adult NMJ phenotypes induced by *peb* knockdown are attributable to a role of Peb in adult axonal maintenance, rather than a role of Peb during development. We first investigated the morphology of the NMJ on muscle 8 of third instar larvae (Figure S4A) with *peb* knockdown in motor neurons. We did not observe any changes in NMJ length (Figure S4B,C), the distribution of the synaptic vesicle protein Synapsin (Greengard *et al*, 1993) (Figure S4D), the SNARE-binding protein Ras opposite (Rop)(Wu *et al*, 1998) (Figure S4E), or the number of active zones (Figure S4F,G). Furthermore, we used a temperature sensitive GAL80 to gain temporal control over *peb* knockdown (Figure 4H), and induced *peb* knockdown in motor neurons either during development (Figure 4I), adulthood (Figure 4J) or both (Figure 4K). Whereas *peb* knock-down during development did not result in tibial NMJ degeneration at 21-days-of-age, knockdown of *peb* during adulthood was sufficient to induce NMJ degeneration, similar to continuous knockdown throughout development and adulthood. Together, these data demonstrate that Peb is required for adult axonal maintenance.

### *peb* mutant axonal degeneration is independent of apoptosis, the JNK signaling pathway and Wallerian degeneration

Next, we explored possible molecular mechanisms by which loss of *peb* function induces axonal degeneration. Peb was shown to be an important negative regulator of Jun kinase (JNK) signaling (Reed *et al*, 2001), a stress-signaling pathway that can induce apoptosis. Given that aberrant activation of cell death pathways is a common feature in neurodegenerative diseases such as Alzheimer’s disease (AD), Parkinson’s disease (PD) and Huntington’s disease (HD) (Moujalled *et al*, 2021), we performed the TUNEL assay to detect apoptosis in homozygous *peb* mutant motor neuron clones in the VNC of 21-day-old flies. No differences in the number of TUNEL positive nuclei compared to control was found (Figure S5A). To exclude the possibility that *peb* mutant motor neurons have died before 21 days of age, we quantified the number of motor neurons in the motor neuron clusters in the VNC that innervate the leg muscles (Phelps *et al*, 2021) (Figure S5B) in 21-day-old animals with *peb* knockdown in motor neurons (*OK371-GAL4*), and found no difference as compared to control (Figure 5A). Expression of a baculovirus anti-apoptotic p35 protein by itself induced NMJ defects in the femur of 21-day-old animals and did not rescue *peb* mutant NMJ morphology defects (Figure 5B). Finally, heterozygosity for a loss-of-function allele of the JNK ortholog *bsk* did not modify *peb* mutant motor axonal and NMJ degeneration (Figure S5C). Together, these data indicate that *peb* mutant motor axonal degeneration is caused by a molecular mechanism independent of apoptosis or the JNK signaling pathway.

**Figure 5:**
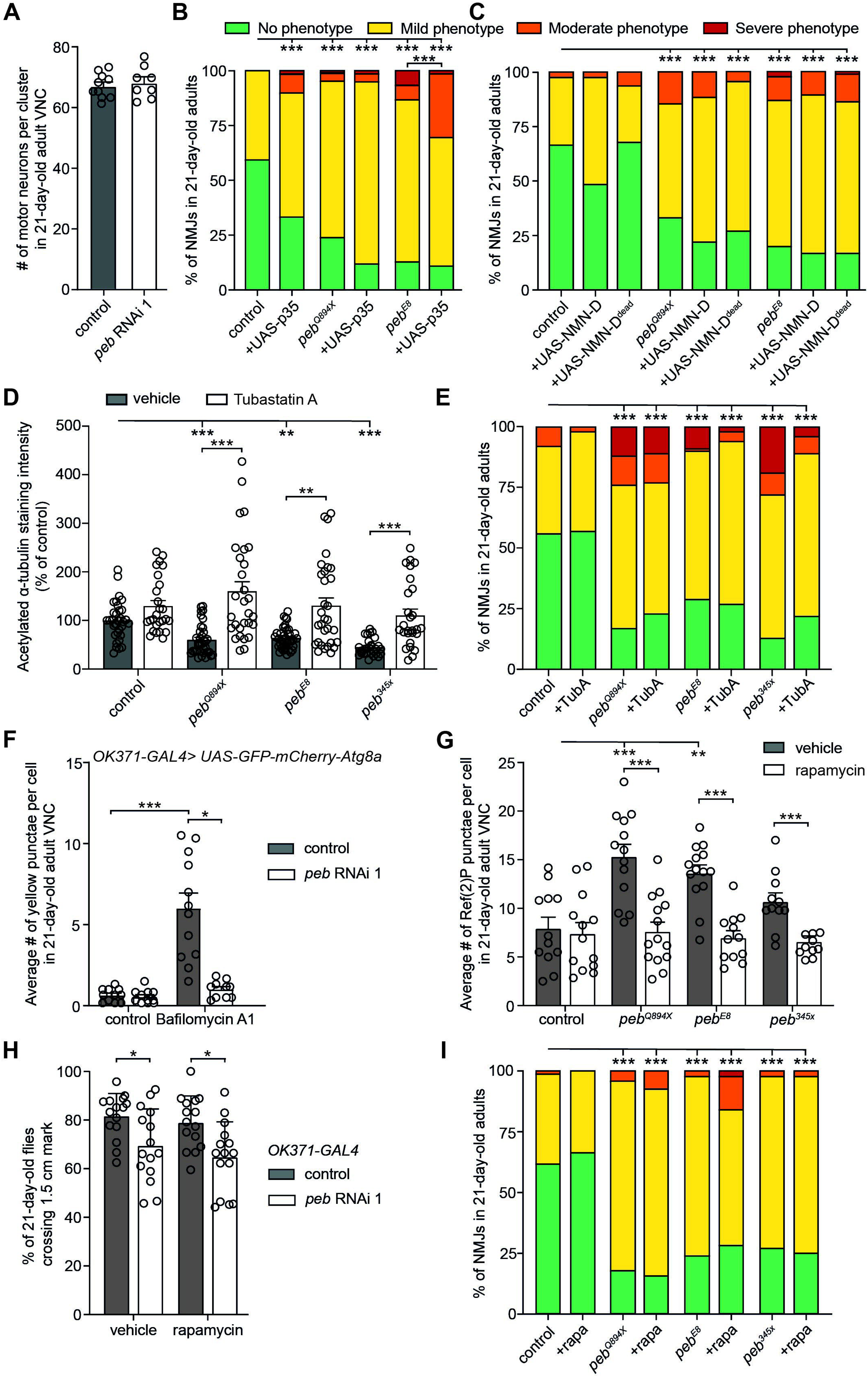
Apoptosis, Wallerian degeneration, defective α-tubulin acetylation or autophagy do not underlie *peb* mutant phenotypes. **(A)** Quantification of the number of motor neurons per cluster in 21-day-old adult VNCs with *peb* knockdown in motor neurons (*OK371-GAL4*) versus driver-only control (KK background). **(B,C)** Semi-quantitative analysis of femoral NMJ degeneration in control (FRT19A) and *peb* mutant motor neurons clones of 21-day-old animals expressing (B) the anti-apoptotic protein p35 or (C) the Wallerian degeneration suppressor NMN-D or NMN-D^dead^. **(D)** Quantification of immunostaining intensity of acetylated α-tubulin in control (FRT19A) versus *peb* mutant motor neuron clones in the VNC of 21-day-old flies treated with 10 μM Tubastatin A or vehicle. Data was normalized to control. **(E)** Quantification of the effect of 10 μM Tubastatin A (TubA) treatment versus vehicle on NMJ degeneration in control (FRT19A) versus *peb* mutant motor neuron clones in the femur of 21-day-old flies. **(F)** Quantification of the effect of *ex vivo* Bafilomycin A1 treatment versus vehicle on the average number of GFP-mCherry-Atg8a yellow punctae per motor neuron (*OK371-GAL4*) in the VNC of 21-day-old control versus *peb* knockdown flies expressing the GFP-mCherry-Atg8a autophagy marker. **(G)** Quantification of the number of Ref(2)P-positive punctae in *peb* mutant versus control (FRT19A) motor neuron clones in the VNC of 21-day-old adult flies treated with either 50 μM rapamycin or vehicle. **(H)** Negative geotaxis motor performance (% of flies crossing a 1.5 cm mark) of 21-day-old flies with *peb* knockdown in motor neurons (*OK371-GAL4*) versus driver-only control (KK background), either treated with vehicle or 50 μM Rapamycin. **(I)** Semi-quantitative analysis of femoral NMJ degeneration in control (FRT19A) and *peb* mutant motor neuron clones of 21-day-old animals treated with either 50 μM rapamycin or vehicle. *P<0.05, **P<0.01, ***P<0.005 by (A) unpaired t-test (B,C,E,I) Fisher’s exact test with Bonferroni correction, (D,F) Kruskal-Wallis test with Dunn’s multiple comparisons test, (G) Brown-Forsythe and Welch ANOVA with Dunnett’s T3 multiple comparisons test, (H) two-way ANOVA with Šídák’s multiple comparisons test. n= (A) 8-10 animals per genotype, (B) 88-116 NMJs in 6-10 animals per genotype, (C) 53-131 NMJs in 6-10 animals per genotype, (D) 8-26 animals per experimental group, (E) 32-182 NMJs of 6-10 animals per experimental group, (F) 6 cells in 10-12 animals per experimental group, (G) 10-14 animals per experimental group, (H) 15 groups of 5-10 animals per experimental group and (I) 54-159 NMJs in 6-10 animals per experimental group. Data are represented as mean ± SEM.

Peb was further reported to be required for Wallerian degeneration, the programmed degeneration of the part of an axon distal to a nerve injury (Coleman & Höke, 2020). Indeed, loss of *peb* function prevents axonal death in wing sensory axons after axotomy (Farley *et al*, 2018). To evaluate the possible involvement of this signaling cascade in *peb* mutant motor axonal degeneration, we generated *peb* mutant motor neuron clones whilst simultaneously altering the levels of dSarm (sterile α/Armadillo/Toll-Interleukin receptor homology domain protein)(Osterloh *et al*, 2012), because this was previously shown to modify *peb* mutant axonal death (Farley *et al*, 2018). Neither heterozygosity for the *dsarm^896^*loss-of-function allele, nor dSarm overexpression did modify the NMJ degenerative phenotype of homozygous *peb^Q894X^* or *peb^E8^* motor neuron clones (Figure S5D,E). Furthermore, we expressed PncC, a bacterial enzyme with NMN-deamidase (NMN-D) activity that can convert the dSarm activating NMN (nicotinamide mononucleotide) to inactive NAMN (nicotinic acid mononucleotide), thus preventing dSarm activation and subsequent degeneration of severed axons (Rosell *et al*, 2022). NMJ degeneration in *peb* homozygous mutant motor neuron clones was not rescued by expression of NMN-D or its enzymatically inactive form, NMN-D^dead^ (Figure 5C). These results demonstrate that the molecular mechanism underlying *peb* mutant motor neuron degeneration does not involve the axon death signaling pathway.

### Defects in α-tubulin acetylation induced by loss of *peb* function are not responsible for *peb* mutant NMJ degeneration

It was previously reported that loss of a nervous system-enriched Rreb1 transcript in mice is associated with decreased α-tubulin glutamylation and acetylation (Griffin *et al*, 2024). Both of these posttranslational modifications are important for microtubule dynamics and axonal transport (Valenstein & Roll-Mecak, 2016; McKenna *et al*, 2023; Janke & Magiera, 2020). Therefore, we investigated α-tubulin glutamylation and acetylation in homozygous *peb* mutant motor neuron clones in 21-day-old animals by quantifying immunostaining intensity. α-tubulin glutamylation was significantly increased in *peb^Q894X^* and *peb^E8^* mutant motor neurons (Figure S5F,G), in contrast to decreased α-tubulin glutamylation in *Rreb1-nm3888^-/-^* mice. However, we found a significant reduction in α-tubulin acetylation in both *peb^Q894X^*and *peb^E8^* homozygous mutant motor neuron clones in the VNC of 21-day-old flies (Figure S6A,B). Microtubule acetylation defects have been linked to several neurodegenerative disorders, including PD (Godena *et al*, 2014), HD (Dompierre *et al*, 2007) and Charcot-Marie-Tooth peripheral neuropathy (d’Ydewalle *et al*, 2011), suggesting that it may causally contribute to *peb* mutant axonal degeneration.

We therefore asked whether selective restoration of α-tubulin acetylation would be sufficient to rescue *peb* loss-of-function phenotypes. We treated flies with Tubastatin A, a frequently used inhibitor of the α-tubulin deacetylase HDAC6 (histone deacetylase 6)(Godena *et al*, 2014; Mao *et al*, 2017; Mo *et al*, 2018), from the adult stage onwards. As expected, Tubastatin A treatment rescued the decrease in α-tubulin acetylation in *peb* homozygous mutant motor neurons at 21 days of age (Figure 5D, representative images in Figure S6C). However, *peb* mutant NMJ phenotypes were not significantly rescued by Tubastatin A treatment (Figure 5E). Thus, rescuing the reduced α-tubulin acetylation induced by loss of *peb* function is not sufficient to rescue the axonal degenerative phenotypes.

### Autophagy defects induced by loss of *peb* function are not responsible for *peb* mutant motor deficits

*Rreb1-nm3888^-/-^* mice also displayed decreased autophagy (Griffin *et al*, 2024) and reduced autophagic clearance has been associated with neurodegeneration in AD, PD and HD (López-Otín *et al*, 2013; Labbadia & Morimoto, 2015; Wilson *et al*, 2023). We therefore evaluated whether defective autophagy might contribute to the neurodegenerative phenotypes induced by loss of *peb* function. Immunostaining for the *Drosophila* p62 ortholog Ref(2)P revealed an increased number of Ref(2)P positive punctae in homozygous *peb* mutant motor neuron clones in 21-day-old flies (Figure S6D,E), indicating accumulation of ubiquitinated proteins. To determine which step of the autophagy pathway is affected, we used an autophagy flux sensor, which is based on the LC3 ortholog Atg8a tagged with GFP and mCherry (Nezis *et al*, 2010; Klionsky *et al*, 2021). This sensor allows to discriminate autophagosomes from autolysosomes, because GFP fluorescence is quenched in the acidic milieu of lysosomes, while mCherry fluorescence persists. Therefore, yellow punctae (positive for both GFP and mCherry) represent autophagosomes, whereas red punctae correspond to autolysosomes (Figure S6F). We observed no changes in the total number of Atg8a-positive punctae (Figure S6G), and no difference in autophagic flux (Figure S6H) in the VNC of 21-day-old flies after *peb* knockdown in motor neurons. To determine whether initiation of autophagy is affected, we performed *ex vivo* Bafilomycin A1 treatment on 21-day-old adult VNCs of flies expressing the autophagy flux sensor with or without knockdown of *peb* in motor neurons (Figure 5F). Bafilomycin A1 inhibits autophagic flux by inhibiting V-ATPase-dependent acidification and preventing autophagosome-lysosome fusion (Mauvezin & Neufeld, 2015). While the number of autophagosomes was not changed in vehicle treated control versus *peb* knockdown motor neurons, *peb* knockdown resulted in significantly decreased accumulation of autophagosomes after Bafilomycin A1 treatment (Figure 5F). This suggests that loss of *peb* function impairs initiation of autophagy, resulting in accumulation of ubiquitinated proteins.

To dissect the possible contribution of decreased autophagy to *peb* mutant phenotypes, we treated flies with rapamycin, an mTORC1 inhibitor which increases autophagy (Bjedov *et al*, 2010; Mannick & Lamming, 2023). Rapamycin treatment fully rescued Ref(2)P accumulation in 21-day-old *peb* mutant motor neuron clones (Figure 5G). Nevertheless, neither the motor performance defect induced by *peb* knockdown in motor neurons, nor the NMJ degeneration induced by loss of *peb* function was rescued by rapamycin treatment (Figure 5H,I). Together, these data indicate that rescuing the autophagy defect is not sufficient to rescue *peb* mutant motor neurodegeneration.

### RAS/MAPK pathway overactivation results in adult-onset motor axonal degeneration

The human Peb ortholog RREB1 was reported to negatively regulate transcription of RAS-MAPK pathway target genes such as *JUN*, *MYC*, *FGFR4*, *HRAS* and *MEK2*, by binding to RAS-responsive elements (RREs) in their promoters (Kent *et al*, 2020). Consistently, *RREB1* haploinsufficiency causes a Noonan-like RASopathy in *Rreb1^+/-^* mice and humans (Kent *et al*, 2020; Shatokhina *et al*, 2024; Strong *et al*, 2025). Indeed, cardiac hypertrophy attributable to excessive RAS/MAPK pathway activation was identified in *Rreb1^+/-^* mice, but neurodegenerative phenotypes have thus far not been reported in these mice (Kent *et al*, 2020). Since RREB1 can functionally replace Peb (Farley *et al*, 2018), they likely function in the same molecular pathways. Furthermore, Peb target genes include several known RAS/MAPK target genes, which are also regulators or effectors of the RAS/MAPK pathway, including *pointed* (*pnt*)(O’Neill *et al*, 1994; Shwartz *et al*, 2013) and *Mkp3* (*Mitogen-activated protein kinase phosphatase 3*)(Butchar *et al*, 2012; Farley *et al*, 2018). We therefore hypothesized that excessive activation of the RAS/MAPK pathway could underlie axonal degeneration in *peb* mutants. Consistent with our hypothesis, RNAi-mediated knockdown of three RAS/MAPK pathway suppressors in motor neurons, *Lztr1*, *Spred*, and *Nf1*(Lorenzo & McCormick, 2020; Bigenzahn *et al*, 2018; Cichowski & Jacks, 2001), resulted in adult-onset NMJ degeneration in the tibia (Figure 6A-C). Consistently, overexpression of a constitutively active form of RAS (Ras^R68Q^) induced a similar phenotype (Figure 6A-C). Moreover, flies heterozygous for a loss-of-function allele of *Neurofibromin 1* (*Nf1*) (Walker *et al*, 2006), a GTPase-Activating Protein (GAP) for RAS that has been associated to NF1, displayed an adult-onset climbing defect at 21 days of age (Figure 6D).

**Figure 6:**
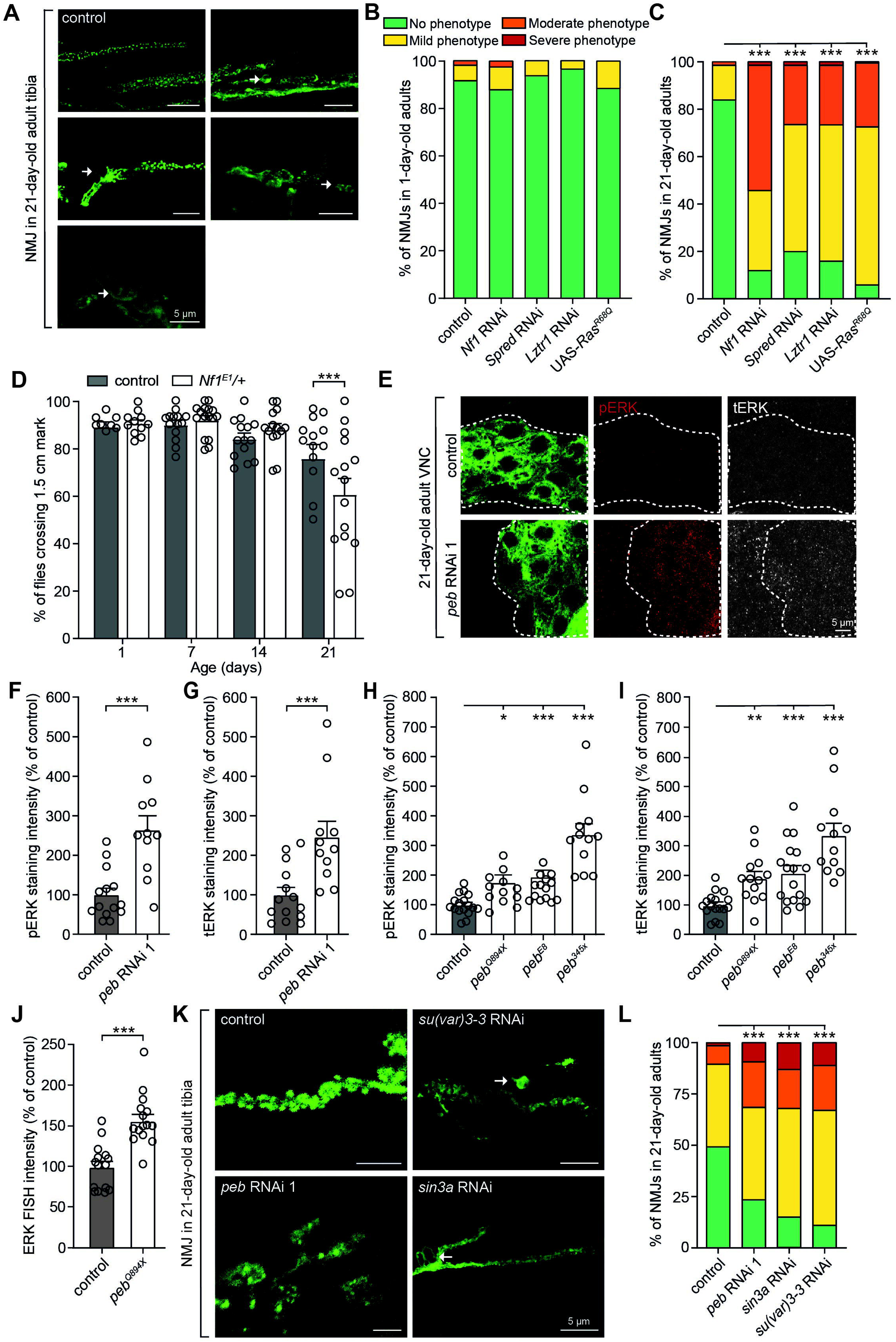
Overactivation of the RAS/MAPK pathway triggers adult-onset NMJ degeneration. **(A)** Representative images and **(B,C)** semi-quantitative analysis of NMJ morphology in the tibia of (B) 1-day-old or (A,C) 21-day-old flies upon knockdown in motor neurons (*OK371-GAL4*) of negative regulators of the RAS/MAPK pathway (Lztr1, Spred, and Nf1), or overexpression of constitutively active Ras (Ras^R68Q^). Arrows indicate NMJ swellings that are typically observed. **(D)** Negative geotaxis motor performance (% of flies crossing a 1.5 cm mark) of heterozygous *Nf1^E1/+^*flies as compared to genetic background control at 1, 7, 14, and 21 days of age. **(E)** Representative images and **(F-I)** quantification of (E,F,H) phospho-ERK (pERK) and (E,G,I) total ERK (tERK) immunostaining intensity in motor neuron cell bodies in the adult VNC at 21 days of age, (E-G) in the presence or absence of *peb* knockdown in motor neurons (*OK371-GAL4*) or (H,I) in *peb* mutant motor neuron clones. Data are normalized to control ((E-G) KK background; (H,I) FRT19A). **(J)** Quantification of signal intensity of FISH for rl (rolled, the *Drosophila* ERK ortholog) transcript in the cell body of *peb^Q894X^* mutant motor neuron clones in the adult VNC at 21 days of age. Data are shown as percentage of control (FRT19A). **(K)** Representative images and **(L)** semi-quantitative analysis of NMJ morphology in the tibia of 21-day-old flies after knockdown of *peb*, *su(var)3-3* or *sin3a* in motor neurons (*OK371-GAL4*). *P<0.05, **P<0.01, ***P<0.005 by (B,C,L) Fisher’s exact test with Bonferroni correction, (D) two-way ANOVA with Šidák’s multiple comparisons test, (F) unpaired t-test with Welch’s correction, (G) Mann-Whitney test, (H,I) Brown-Forsythe and Welch’s ANOVA with Dunnett’s T3 test for multiple comparisons, and (J) unpaired t-test. n= (B) 99-162 NMJs in 6-10 animals per genotype, (C) 88-131 NMJs in 6-10 animals per genotype, (D) 8-16 groups of 10 flies per genotype, (F,G) 11-14 animals per genotype, (H,I) 12-17 animals per genotype, (J) 15 animals per genotype, and (L) 139-204 NMJs of 6-10 animals per genotype. Data are represented as mean ± SEM. Scale bars in A,E,K: 5 μm.

To biochemically confirm RAS/MAPK pathway overactivation upon loss of *peb* function, we used phospo-ERK (pERK) and total-ERK (tERK) antibodies (Botero *et al*, 2021) to quantify the levels of ERK ortholog Rolled in motor neuron cell bodies in 21-day-old adult VNCs after *peb* knockdown or in *peb* mutant clones (Figure 6E-I). We found a significantly increased staining intensity of both pERK (Figure 6E,F,H) and tERK (Figure 6E,G,I) following *peb* knock-down and in all three *peb* mutants. The increased tERK levels induced by loss of *peb* function suggest that Peb negatively regulates *rolled* transcription. Consistently, *rolled* mRNA levels are increased in cell bodies of 21-day-old *peb^Q894X^* motor neuron clones (Figure 6J). Taken together, our data show that RAS/MAPK pathway overactivation, in *peb* mutant conditions and otherwise, results in adult-onset motor neurodegeneration.

After binding to RREs, RREB1 recruits SIN3A (SIN3 Transcription regulator Family Member A) and KDM1A (Lysine Demethylase 1A) to regulate histone H3K4 methylation at RAS/MAPK target gene promoters and reduce their expression (Kent *et al*, 2020). Similarly, Rreb1 forms a complex with Sin3a and Kdm1a in mice (Kent *et al*, 2020). We therefore evaluated the effects of knockdown of *sin3a* and *su(var)3-3*, the *Drosophila* orthologs of *SIN3A* and *KDM1A*, respectively. Similar to *peb* loss-of-function, knockdown of either *sin3a* or *su(var)3-3* in motor neurons induced an NMJ degenerative phenotype in 21-day-old adults (Figure 6K,L). These data indicate that in adult motor neurons, Peb likely functions in a complex with Sin3a and Su(var)3-3 to negatively regulate the transcription of RAS/MAPK pathway target genes in *Drosophila*.

### Pharmacological or genetic inhibition of the RAS/MAPK pathway rescues *peb* and *Nf1* loss-of-function phenotypes

To confirm that excessive activation of the RAS/MAPK pathway triggers adult-onset axonal degeneration in *peb* mutants, we evaluated the effect of pharmacological inhibition of the RAS/MAPK pathway on *peb* loss-of-function phenotypes. We treated adult flies with *peb* knockdown in motor neurons (*OK371-GAL4*) with mirdametinib, an FDA-approved small molecule MEK1/2 inhibitor that is highly blood-brain-barrier penetrable (Weiss *et al*, 2021; Moertel *et al*, 2024; Vinitsky *et al*, 2022). We observed a dosage-dependent rescue of the motor performance deficit in 21-day-old *peb* knockdown animals (Figure 7A), with 0.1 µM mirdametinib resulting in partial rescue and 1 µM mirdametinib inducing full rescue of the motor deficit. Furthermore, 1 µM mirdametinib fully rescued the NMJ degenerative phenotype of *peb* mutant motor neuron clones (Figure 7B,C). To confirm that the phenotypic rescue induced by mirdametinib treatment is attributable to inhibition of the *Drosophila* MEK1/2 ortholog *Dsor1*, we knocked down *Dsor1* in *peb* mutant motor neuron clones. As expected, *Dsor1* knockdown rescued NMJ degeneration induced by loss of *peb* function (Figure 7D). Importantly, NMJ degeneration in 21-day-old *Nf1^E1/+^* heterozygous flies or flies with *Nf1* knockdown in motor neurons was also fully rescued by 1 µM mirdametinib (Figure 7E). Consistently, mirdametinib treatment rescued the increased levels of phospho-ERK in the VNC of 21-day-old *Nf1^E1/+^*heterozygous flies (Figure 7F). Together, these data confirm that motor neurodegenerative phenotypes induced by loss of *peb* or *Nf1* function are attributable to overactivation of the RAS/MAPK pathway.

**Figure 7:**
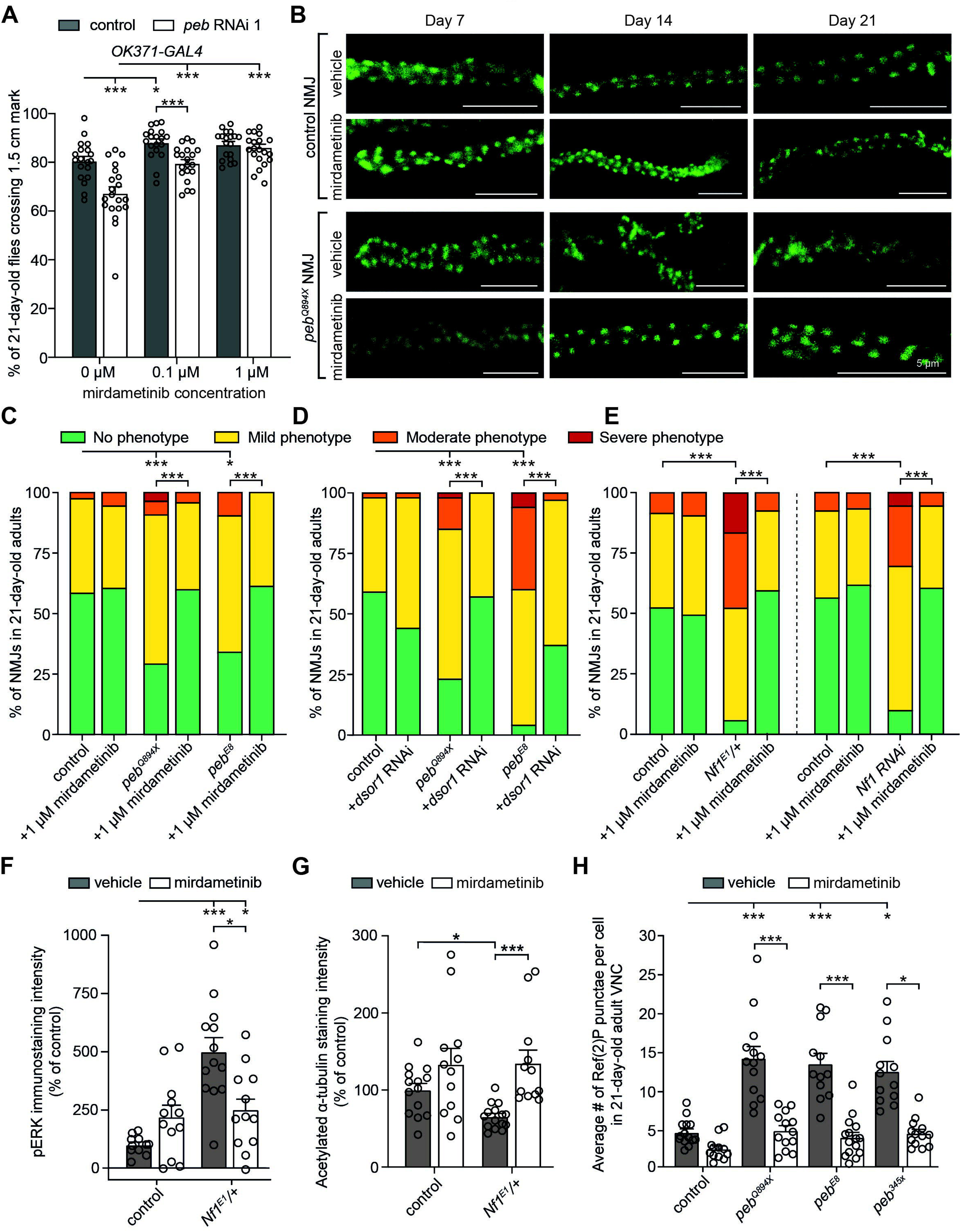
Pharmacological or genetic inhibition of the RAS/MAPK pathway rescues *peb* and *Nf1* loss-of-function phenotypes. **(A)** Negative geotaxis motor performance of 21-day-old flies with *peb* knockdown in motor neurons (*OK371-GAL4*) versus driver-only control (KK-background), treated with vehicle, 0.1 μM or 1 μM mirdametinib. **(B)** representative images and **(C)** semi-quantitative analysis of the effect of 1μM mirdametinib versus vehicle treatment on NMJ degeneration in *peb* mutant versus control (FRT19A) motor neuron clones in the femur of 21-day-old flies. **(D)** Semi-quantitative analysis of the effect of *Dsor1* knockdown on NMJ degeneration in *peb* mutant versus control (FRT19A) motor neuron clones in the femur of 21-day-old flies. **(E)** Semi-quantitative analysis of the effect of 1μM mirdametinib treatment versus vehicle on NMJ degeneration in the tibia of 21-day-old heterozygous *Nf1^E1^/+* flies or flies with *Nf1* knockdown in motor neurons (*OK371-GAL4*) versus genetic background controls. **(F)** Quantification of phospho-ERK (pERK) immunostaining intensity in motor neuron cell bodies in the adult VNC of 21-day-old heterozygous *Nf1^E1/+^* versus control flies, either treated with 1μM mirdametinib or vehicle. Data are normalized to control. **(G)** Quantification of immunostaining intensity of acetylated α-tubulin in motor neurons of 21-day-old heterozygous *Nf1^E1/+^* versus control flies, either treated with 1μM mirdametinib or vehicle. Data were normalized to control. **(H)** Quantification of Ref(2)P-positive puncta in *peb* mutant versus control (FRT19A) motor neuron clones in the VNC of 21-day-old flies, either treated with 1μM mirdametinib or vehicle. *P<0.05, ***P<0.005 by (A,F,G) Brown-Forsythe and Welch’s ANOVA with Dunnett’s T3 test for multiple comparisons, (C,D,E) Fisher’s exact test with Bonferroni correction or (G,H) Kruskal-Wallis test with Dunn’s multiple comparisons test. n= (A) 19-20 groups of 10 flies per experimental group, (C) 66-100 NMJs of 6-10 animals per experimental group, (D) 43-101 NMJs in 6-10 animals per experimental group, (E) 96-140 NMJs of 6-10 animals per experimental group, (F) 11-13 animals per experimental group, (G) 12-15 animals per experimental group and (H) 12-15 animals per experimental group. Data are represented as mean ± SEM. Scale bars: 5 μm. See also Figure S7.

### Defects in α-tubulin acetylation and autophagy are downstream of RAS/MAPK pathway overactivation

The RAS/MAPK pathway has many downstream targets that may possibly be involved in neuronal maintenance (Mendoza *et al*, 2011; Deng *et al*, 2020), including HDAC6 (Williams *et al*, 2013) and mTOR (Mendoza *et al*, 2011). To determine whether the reduced α-tubulin acetylation levels and decreased autophagy are a consequence of RAS/MAPK pathway overactivation, we treated flies with mirdametinib or vehicle and evaluated acetylated α-tubulin levels and Ref(2)P-positive punctae at 21 days of age. Mirdametinib treatment restored acetylated α-tubulin levels in *peb^Q894X^* mutant motor neuron clones (Figure S7A,B) and in heterozygous *Nf1^E1/+^* animals (Figure 7G). Furthermore, increased Ref(2)P accumulation in *peb* mutant motor neuron clones was fully rescued by mirdametinib treatment (Figure 7H, representative images in Figure S7C), thus demonstrating that the reduced α-tubulin acetylation levels and decreased initiation of autophagy are downstream of excessive RAS/MAPK signaling.

## Discussion

In this study, we performed a genetic screen to identify genes on the *Drosophila* X chromosome that are required for maintenance of peripheral motor axons and NMJs. The deployed screening approach is unique because it utilizes microscopy imaging to identify neurodegenerative changes in motor and sensory neurons, their axons and NMJs in legs of 21-day-old flies. This phenotypic read-out had not previously been used in *Drosophila* forward genetic screens. We identified mutations in 30 genes, of which 29 (97%) have human orthologs and 19 (66%) have previously been associated with neurological disorders, underscoring the disease-relevance of our screen. For these neurological disease-associated genes, the identified mutants may provide a *Drosophila* model for the respective diseases. For three identified genes (*Arp8*, *CG15890* and *Tim9a*), including one disease-associated gene, no hemizygous lethal loss-of-function alleles were available until now.

10 (34%) of the 29 identified genes with human orthologs have not yet been associated with neurological disease. Four of these genes (*Arp10*, *sno, Idh3a* and *AMPKα*) have been linked to neurological phenotypes in animal models. For example, loss of function of the zebrafish *Arp10* ortholog *actr10* results in defective retrograde axonal transport of mitochondria and abnormal axon morphology (Mandal *et al*, 2021; Drerup *et al*, 2017), whereas cortical neuron-specific knockout of the *sno* ortholog *Sbno1* in mice resulted in hypotrophic axon bundles (Erkhembaatar *et al*, 2022). Four other genes (*Arp8*, *Tim9a*, *sas10* and *Hr4*) were reported to function in a pathway that was previously linked to neurodegeneration, but thus far, no neuronal phenotypes had been reported for these specific genes. For example, Arp8 ortholog ACTR8 has been shown to be a subunit of the INO80 chromatin remodeling complex, which plays an important role in DNA damage repair (Kashiwaba *et al*, 2010). Defective DNA damage repair has been extensively studied as a cause for neurodegeneration (Madabhushi *et al*, 2014), however, ACTR8 loss-of-function has never before been directly linked to neurological phenotypes. Finally, for *Tsp2a*, a tetraspanin that localizes to smooth septate junctions in the *Drosophila* midgut (Izumi *et al*, 2016), no neuronal functions had been described thus far. Thus, our genetic screen uncovered for the first time a neuronal function for five of the identified genes.

The fact that the majority of identified axonal degeneration genes have human orthologs associated with human neurodegenerative or neurodevelopmental disease suggests that the 10 identified genes that have not yet been linked to human neurological disease may represent candidate disease genes. Indeed, while a fraction of recessively inherited or X-linked neurodegenerative and neurodevelopmental disease genes has been identified, more genes remain to be identified, because mutations in the causative genes may be rare, large families with multiple affected members may not be available, and human populations are ‘outbred’, with every individual carrying numerous potentially disease-causing variants. Thus, the human orthologs of the identified genes may help prioritize candidate genes located in linked homozygous regions of human neurodegenerative and neurodevelopmental disease patients with a recessive inheritance pattern. Remarkably, even though our screening design was tailored to identify genes that cause neurodegeneration through a (partial) loss-of-function mechanism, 63% (22 out of 35) of the neurological disorders associated with the human orthologs of identified genes display an autosomal dominant inheritance pattern. This implies that the human orthologs of the 10 *Drosophila* genes that have hitherto not been associated with neurological disease may represent candidate genes for both recessive and dominantly inherited neurological disorders. Furthermore, the human orthologs of identified genes may constitute candidate (polygenic) risk genes for sporadic forms of human neurodegenerative diseases, e.g. in case they are located in a genomic region linked to a risk allele identified in GWAS studies (Diaz-Ortiz & Chen-Plotkin, 2020; Lambert *et al*, 2023). Thus, the genes identified in our screen could help prioritize functional investigation of GWAS hits.

Remarkably, we identified mutations in 12 *Drosophila* genes with human orthologs associated with neurodevelopmental disorders that induce neurodegenerative phenotypes in *Drosophila*. On the one hand, altered neurodevelopmental processes might sensitize neurons to become more susceptible to degeneration with ageing (Hickman *et al*, 2022; Van Battum *et al*, 2015). On the other hand, proteins implicated in neurodegenerative disorders often play important roles in brain development, opening the possibility of shared pathogenic mechanisms (Schor & Bianchi, 2021). Indeed, for neurogenetic conditions associated with intellectual disability and autism spectrum disorders, emerging natural history data indicate that older adult patients may develop neurodegeneration. For instance, Down syndrome (trisomy 21) patients develop neuropathological hallmarks of AD by 40 years of age, and subsequently often develop dementia (Hickman *et al*, 2022). Currently, only a few of the identified genes that are associated with neurodevelopmental disorders have previously been associated with neurodegenerative or neuropathy phenotypes. One example is the *ý-Spec* ortholog *SPTBN4*, which is associated with NEDHND, a neurodevelopmental disorder with hypotonia, motor axonal neuropathy, and deafness (Wang *et al*, 2018; Häusler *et al*, 2020). Our data suggests that other neurodevelopmental disorders may also be associated with axonal degeneration. More research is needed to identify possible neurodegenerative features in individuals with neurodevelopmental disorders.

Taken together, our genetic screen provides important novel insight into the molecular mechanisms underlying axonal degeneration. Importantly, transgenic knock-down of identified axonal degeneration genes revealed that only a small minority of genes would have been identified in a transgenic RNAi screen. The most likely explanation is that RNAi-mediated knock-down is incomplete and induces only partial loss-of-function, whereas EMS-induced mutations may yield full loss-of-function alleles. This underscores the value of EMS-based genetic screens and illustrates the complementarity of EMS and RNAi-based genetic screens. We believe that it would be valuable to apply the same screening approach to the second and third chromosomes, which comprise about 80% of the *Drosophila* genome.

One of the identified axonal degeneration genes is *pebbled* (*peb*), the *Drosophila* ortholog of human *RREB1*. Loss of *peb* function resulted in adult-onset degeneration of motor axons and NMJs, as well as motor performance deficits. These defects are mediated by excessive RAS/MAPK pathway activity. Peb expression is highly dynamic, and Peb mediates various functions depending on the specific cell type and life cycle stage (Deady *et al*, 2017; Baechler *et al*, 2015). For example, Peb was reported to positively regulate EGFR signaling and to downregulate JNK signaling in *Drosophila* embryos (Kim *et al*, 2020; Melani *et al*, 2008; Reed *et al*, 2001). In contrast, in adult motor neurons, we find that *peb* mutant phenotypes are not mediated by JNK-induced apoptosis. In addition, loss of *peb* function in adult motor neurons results in progressive degeneration that is unrelated to Wallerian degeneration, whereas in sensory neurons, *peb* loss-of-function does not induce degeneration (this study), and prevents Wallerian degeneration (Farley *et al*, 2018). The wide variety of Peb functions in different cell types may be attributable to epigenetic regulation of chromatin accessibility and the expression pattern of Peb-interacting proteins in a given cell type and life cycle stage. Indeed, Peb may regulate a different set of target genes depending on chromatin accessibility and the protein complexes that Peb associates with. For instance, RREB1 has been shown to interact with several chromatin remodeling and histone modification complexes, such as HDAC1, HDAC2, KDM2A, and KDM3B (Kent *et al*, 2020).

Our data suggest that overactivation of the RAS/MAPK pathway may be involved in neurodegenerative disease pathogenesis. Consistent with this notion, excessive RAS/MAPK pathway activation was previously implicated in *LRRK2*-associated PD. Indeed, in patient iPSC-derived dopaminergic neurons, the LRRK2 G2019S mutation was associated with increased ERK1/2 phosphorylation, and mirdametinib mitigated several PD-associated phenotypes in this cellular model (Reinhardt *et al*, 2013). Furthermore, increased ERK1/2 phosphorylation has been associated with AD (Pei *et al*, 2002) and ALS (Ayala *et al*, 2011b), and multiomic analysis suggested the MAPK pathway as an early disease mechanism in ALS (Caldi Gomes *et al*, 2024). However, whether increased phospho-ERK1/2 levels causally contribute to neurodegeneration in humans remains to be investigated. Our data are the first to show that increased RAS/MAPK signaling is sufficient to induce adult-onset neurodegeneration. Interestingly, loss of a nervous system-enriched *Rreb1* transcript (*V7* isoform) in mice induces progressive loss of cerebellar Purkinje cells (Griffin *et al*, 2024), and in knock-in mouse models of Legius syndrome, a heterozygous P415A or P415V mutation in *Spred1* resulted in late-onset Purkinje cell degeneration leading to cerebellar ataxia (Hirata *et al*, 2024). Our findings suggest that these phenotypes may be attributable to excessive activation of the RAS/MAPK pathway.

What may be the molecular mechanism downstream of excessive RAS/MAPK pathway activation that mediates axonal degeneration? While we could show that autophagy is impaired and α-tubulin acetylation is reduced in *peb* mutant motor neurons as a consequence of RAS/MAPK pathway overactivation, pharmacological rescue of these pathways individually was not sufficient to prevent axonal degeneration. Possibly, dysregulation of each of these pathways – and possibly additional ones – is sufficient to induce neurodegeneration, which would explain why rescuing one downstream pathway alone does not rescue *peb* mutant neurodegeneration. Alternatively, another molecular mechanism downstream of increased RAS/MAPK pathway activation, unrelated to autophagy or α-tubulin acetylation, may mediate *peb* mutant axonal degeneration. More work is needed to unravel the precise molecular mechanisms responsible for neurodegeneration downstream of RAS/MAPK pathway overactivation.

Our finding that loss of function of several negative regulators of the RAS/MAPK pathway induced motor axonal and NMJ degeneration suggests that neurodegenerative phenotypes may represent a currently underappreciated feature of RASopathies. In particular, our findings that heterozygous loss of *peb* or *Nf1* function is sufficient to induce age-dependent motor deficits in *Drosophila* suggests that haploinsufficiency of these genes – as is the case in *RREB1*-associated Noonan-like syndrome and NF1 – is sufficient to induce axonal degeneration. Consistently, beyond cerebellar Purkinje cell degeneration in *RREB1*-mutant mice (Griffin *et al*, 2024) and mouse models of Legius syndrome (Hirata *et al*, 2024), *Nf1^+/-^* mice display motor deficits that are not attributable to cerebellar dysfunction (van der Vaart *et al*, 2011).

Interestingly, progressive neuropathic pain and muscle weakness associated with nerve enlargement were reported in children and adults with Noonan syndrome (De Ridder *et al*, 2022; Draaisma *et al*, 2023, 2024). Furthermore, a subset of NF1 patients displays neurofibromatous neuropathy, which is typically associated with large numbers of neurofibromas – the hallmark lesion of NF1 – and disruption of the perineum (Ferner *et al*, 2004; Drouet *et al*, 2004). However, beyond tumor-dependent neuropathies, tumor-independent neuropathies affect ∼2% of NF1 patients (Schulz *et al*, 2018). While the mechanism underlying neuropathy phenotypes in Noonan syndrome and related RASopathies remains unclear, our findings suggest that enhanced RAS/MAPK signaling in motor neurons may contribute to neuropathy phenotypes. In the future, this could be explored in clinical research and in RASopathy mouse models. Given that we could fully rescue all motor deficits by mirdametinib treatment in *Drosophila* RASopathy models, MEK1/2 inhibition might serve as a potential therapeutic approach, not only for Noonan syndrome, but more broadly for RASopathy patients that suffer from neuropathies, and potentially also for patients suffering neurodegenerative diseases.

## Methods

### Reagents and Tools Table

| Reagent/Resource | Reference or Source | Identifier or Catalog Number |
| --- | --- | --- |
| **Experimental Models** |  |  |
| *Drosophila melanogaster* | See Experimental Model and Subject Details and Data S4 |  |
| **Antibodies** |  |  |
| Mouse monoclonal anti-Dlg1 | DSHB | Cat#: 4F3,<br>RRID: AB_528203 |
| Mouse monoclonal anti-Brp | DSHB | Cat#: Nc82<br>RRID: AB_2314866 |
| Mouse monoclonal anti-Syn | DSHB | Cat#: 3C11<br>RRID: AB_528479 |
| Mouse monoclonal anti-Rop | DSHB | Cat#: 4F8<br>RRID: AB_1157869 |
| Mouse monoclonal anti-Peb | DSHB | Cat#: 1G9<br>RRID: AB_528278 |
| Rabbit polyclonal anti-Ref2P | Abcam | Cat#: Ab178440<br>RRID: AB_2938801 |
| Mouse monoclonal anti-polyglutamylated alpha-tubulin, Clone GT355 | Avantor | Cat#: # USBI045161 |
| Mouse monoclonal anti-acetylated alpha-tubulin, Clone 6-11B-1 | Sigma Aldrich | Cat#: MABT868<br>RRID: AB_2819178 |
| Mouse monoclonal anti-phospho-ERK | Sigma Aldrich | Cat#: M8159<br>RRID: AB_477245 |
| Rabbit polyclonal anti-ERK | Cell Signaling | Cat#: 9102 |
| Rabbit anti-GFP | Invitrogen | Cat#: A6455<br>RRID: AB_221570 |
| Alexa Fluor goat anti-mouse-568 | Invitrogen | Cat#: A11004<br>RRID: AB_2534072 |
| Alexa Fluor Goat anti-rabbit-568 | Invitrogen | Cat#: A11011<br>RRID: AB_143157 |
| Alexa Fluor goat anti-rabbit-647 | Invitrogen | Cat#: A21245<br>RRID: AB_2535813 |
| Alexa Fluor Goat anti-mouse-647 | Invitrogen | Cat#: A21235<br>RRID: AB_2535804 |
| Alexa Fluor Goat anti-rabbit-488 | Invitrogen | Cat#: A11008<br>RRID: AB_143165 |
| **Oligonucleotides and other sequence-based reagents** |  |  |
| Sanger sequencing primers | See data S4 |  |
| **Chemicals, Enzymes and other reagents** |  |  |
| EMS | Sigma-Aldrich | M0880; CAS: 62-50-0 |
| Sucrose | Sigma-Aldrich | S9378; CAS: 57-50-1 |
| RNase A | Thermo Fisher | EN0531 |
| Bovine Serum Albumin | Sigma-Aldrich | A9647 |
| Formaldehyde solution | Sigma-Aldrich | 252549 |
| Bouin's solution | Sigma-Aldrich | HT10132-1L |
| Goat serum | Gibco | 16210064 |
| Mirdametinib | MedChemExpress | HY-10254G |
| Rapamycin | MedChemExpress | HY-10219 |
| Bafilomycin A1 | MedChemExpress | HY-100558 |
| Tubastatin-A | MedChemExpress | HY-13271A |
| Vectashield Antifade Mounting Medium | Vector Labs | H1000-10 |
| Prolong Glass antifade Mountant | Thermo Fisher | P36980 |
| Triton-X-100 | Sigma-Aldrich | T8787; CAS:9036-19-5 |
| Tween 20 | Sigma-Aldrich | P1379 ; CAS: 9005-64-5 |
| Dimethyl Sulfoxide (DMSO) | Sigma-Aldrich | D2438 |
| Ultrapure SSC, 20X | Invitrogen | 15557044 |
| Sodium Citrate Dihydrate | Fisher Scientific | S279-500 |
| 100% ethanol | VWR Chemicals | 20821.296; CAS:64-17-5 |
| DNase RNase free 100% ethanol | Fisher Bioreagents | BP2818-500; CAS: 64-17-5 |
| Liquid Proteinase K | Macherey-Nagel | 740396 |
| Nuclease-free water | Invitrogen | 10977-035 |
| Methanol | Fisher Chemical | M/4000/17; CAS:67-56-1 |
| Jazz-Mix | Fisher Scientific | AS153 |
| Critical commercial assays |  |  |
| DNeasy Blood & Tissue kit | Qiagen | 69504 |
| Illumina Nextera Flex | Illumina | 20018704 |
| Illumina Nextera CD Indexes | Illumina | 20018707 |
| NextSeq 500/550 High Output Kit v2.5 (300 Cycles) | Illumina | 20024908 |
| In Situ Cell death detection kit, TMR red | Roche | 12156792910 |
| Qubit 1x dsDNA HS Assay Kit | Thermo Fisher | Q33231 |
| Agilent DNA 12000 Kit | Agilent | 5067-1508 |
| Phusion Plus DNA polymerase | Thermo Fisher | F630L |
| FISH kit | Molecular instruments | N/A |
| GeneJET Gel Extraction and DNA cleanup Micro Kit | Thermo Scientific | K0832 |
| **Software** |  |  |
| GraphPad prism V10.2.1 |  |  |
| Image J | (Schindelin *et al*, 2012) | https://imagej.net/software/fiji/ |
| Deadeasy synapse | (Sutcliffe *et al*, 2013) | https://www.biosciences-labs.bham.ac.uk/hidalgo/Software.html |
| FastQC v0.11.5 | (Andrews, 2010) | https://www.bioinformatics.babraham.ac.uk/projects/fastqc/ |
| Cutadapt v1.9.1 | (Martin, 2011) | https://cutadapt.readthedocs.io/en/stable/ |
| Samtools v1.11 | (Li *et al*, 2009) | https://www.htslib.org/ |
| Picard v2.25.4 | Broad Institute | https://broadinstitute.github.io/picard/ |
| BWA v0.7.12-r1039 | (Li, 2013) | https://github.com/lh3/bwa |
| Gatk v4.2.0.0 | (Poplin *et al*, 2018) | https://gatk.broadinstitute.org/hc/en-us |
| Snpeff v5.0 | (Cingolani *et al*, 2012) | https://pcingola.github.io/SnpEff/ |
| Snpsift v5.0 | (Ruden *et al*, 2012) | https://pcingola.github.io/SnpEff/#snpsift |

## Methods and protocols

### Drosophila husbandry

Flies were maintained on standard fly food at 25°C unless otherwise described. Table S1 lists mutants generated in this study. Duplication lines used for mapping, null alleles for complementation crosses, RNAi lines and other *Drosophila* genotypes are described in Table S4. Age and sex of flies (*Drosophila melanogaster*) used in this study, including experimental crosses, are described in Table S5.

### Isogenization and mutagenesis

Isogenized FRT19A/Y males were starved for 8 h before being treated with 10mM EMS (M0880, Sigma-Aldrich) in 1% sucrose (S9378, Sigma-Aldrich) solution on Whatmann paper for 16 h (Bökel, 2008). After 8 h of recovery, EMS-treated males were crossed to *caz^2^*/Fm7i virgins. Parents were removed from the cross after 3 days. See Figure S1 for crossing scheme. Non-hemizygous lethal stocks were discarded. Saturation was calculated by *S* = p(N≥1) = 1 - e ^-(n/E^ * ^λ)^, where S is the saturation (Bökel, 2008). E equals the number of essential genes on the X chromosome, which is estimated to be around 690. n equals the number of genomes screened, thus 8892. λ can be calculated by the probability of receiving 0 lethal hits, P(N=0). This probability equals 0.84075 (8892 total stocks/ 7476 stocks that have not received a lethal hit). This results in a λ of 0.17. Thus, S=1-e ^-(8892/690^ * ^0.17)^ =0.88.

### In vivo leg imaging & screening

#### Initial MARCM screening

6 legs of at least two 21-day-old females were dissected and mounted in ProLong Glass antifade mountant (P36980, Thermo Scientific) and examined for motor axon and NMJ integrity under a fluorescence microscope (DM2500, Leica) as previously described (Sreedharan *et al*, 2015). If a phenotype was observed, crosses were repeated to confirm the phenotype. If no GFP-labelled neurons were observed after three independent crosses, stocks were marked cell-lethal.

#### Confocal microscopy

For time course screening or phenotypic characterization of phenotypes in adult legs, 6 legs of at least 6 flies were examined and imaged on a Leica Sp8 Liachroic confocal microscope using a 63x objective. Axon images were taken from mid-femur or proximal tibia. NMJ images were taken from proximal femur or mid-tibia.

#### Semi-quantitative analysis

To quantify NMJ phenotypes, all legs of 6-10 flies were imaged by confocal microscopy (63x objective, 0.75x zoom). NMJs were scored as “no phenotype”, “mild phenotype”, “moderate phenotype” or “severe phenotype” based on the following criteria (representative images shown in Figure 3C):

|  |  | **Normal** | **Mild** | **Moderate** | **Severe** |
| --- | --- | --- | --- | --- | --- |
| NMJ | Active zones | Aligned and consecutive | Partial loss of alignment | Completely misaligned | Not recognizable |
|  |  | Round | Slightly distorted | Enlarged GFP punctae |  |
|  | Shape | Straight | Slightly deformed | Severely deformed, membrane swellings | Complete loss of structure |
|  |  | Elongated |  | Reduced NMJ length | Almost fully degenerated |

### Duplication and complementation mapping

All mutants that showed a motor axonal phenotype were subsequently crossed to the (1:Y) duplication stocks from the Bloomington Duplication Project (Cook *et al*, 2010) (see Table S4 for all stocks used). Once rescued males were found, stocks were crossed to 1;3 linked P[acman] BAC duplications within the rescued regions. If no (1;Y) rescue was observed, stocks were crossed to 1;3 P[acman] BAC duplications covering regions that were not covered by (1;Y) duplications. All rescued stocks were crossed *inter se* with other mutants rescued by the same duplication to establish complementation groups. Additionally, rescued mutant males were crossed to all non-rescued and cell-lethal mutants for complementation mapping (see Figure S2).

### Genome isolation

Genomes of heterozygous (FRT19A*/FRT19A) and transheterozygous (FRT19A*^1^/FRT19A*^2^) mutants were isolated, together with the isogenized FRT19A. 50 virgins were collected and starved for 8 hours on Whatmann paper with water. Flies were flash-frozen and crushed using mortar and pestle. DNeasy Blood & Tissue Kit (69504, Qiagen) was used for subsequent genomic isolation. 180 μl of buffer ATL and 20 μl of ProtK was added to the crushed flies and incubated overnight at 56 °C. ProtK was heat-inactivated for 20 min at 75°C. 20 μL RNAse A (EN0531, Thermo Scientific) was added and samples were incubated for 2h at 37°C. 200 μl buffer AL and 200 μl DNAse-RNAse-free 100% ethanol (BP2818, Fisher Bioreagents) were then added and mixed. The mixture was pipetted with a cut pipet tip onto a spin column and centrifuged at 6000 x g for 1 min. 500 μl buffer AW1 was added before centrifuging again at 6000 x g for 1 min. 500 μl buffer AW2 was added and centrifuged at 20.000 x g for 3 min. The column was air-dried for several minutes. 50 μl of preheated nuclease free water (10977-035, Invitrogen) was added and incubated for 15 min at 70°C. DNA was eluted by centrifugation for 1 min at 6.000 x g. The elution step was repeated to obtain a higher DNA yield. Nanodrop, Qubit (Q33231, Thermo Fisher) and agarose gel were used to assess quality and quantity of DNA.

### Library prep and whole-genome sequencing

All mutants that showed a motor axonal phenotype, as well as non-complementing cell-lethal mutants were used for whole-genome sequencing (See also Figure S2). Mutants were sequenced transheterozygously as described (Haelterman *et al*, 2014). Illumina Nextera Flex (20018704, Illumina) was used for library preparations together with Nextera CD Indexes (20018707, Illumina) according to manufacturer’s instructions. Library concentrations were quantified using Qubit and fragment size was estimated by Bioanalyzer (5067-1508, Agilent). Libraries were loaded on a 300 cycle High Output flowcell (20024908, Illumina) and sequenced on a NextSeq 500.

### Whole genome sequencing data analysis

To assess the quality of Illumina sequencing reads, FastQC (version 0.11.5) was used (Andrews, 2010). Cutadapt (version 1.9.1)(Martin, 2011) was then used to trim adapter sequences wherever necessary. Alignment against the reference genome (*Drosophila melanogaster* release 6.32 plus ISO1 MT, Refseq: GCF_000001215.4) was performed using Burrows-Wheeler Aligner software (BWA version 0.7.12-r1039) using the BWA-MEM algorithm (Li, 2013). Samtools (version 1.11)(Li *et al*, 2009) was used for sorting and indexing of the aligned reads. Duplicates were removed using Picard (version 2.25.4) MarkDuplicates. Genome Analysis Toolkit (GATK version 4.2.0.0) was used for base quality recalibration and subsequent variant calling using HaplotypeCaller (Poplin *et al*, 2018). SnpEff (version 5.0)(Cingolani *et al*, 2012) and SnpSift (version 5.0)(Ruden *et al*, 2012) were used for variant annotation and filtering respectively.

For rescued mutants, genomic coordinates of the 1;3 P[acman] duplications were used for filtering purposes. For non-rescued mutants, variants were filtered for nonsense single nucleotide variances (SNVs) or INDELS over the entire X-chromosome. Only heterozygous SNVs or INDELS with quality score >25 were considered. All identified SNVs were compared to the isogenized FRT19A genome and all other mutant genomes to remove any pre-existing, non-disease-causing variants. Human orthologs were identified through Ensembl (Harrison *et al*, 2024), MARRVEL (Wang *et al*, 2017) and Alliance (Alliance of Genome Resources Consortium, 2024).

### Phenotypic rescue

1;3 P[acman] duplication lines of all candidate disease-causing variants were introduced on the third chromosome of each mutant to determine 1) whether the lethality was rescued and 2) whether the axonal/NMJ degeneration phenotype was rescued (See Table S4 for the duplication lines used). To assess the phenotypic rescue, MARCM crosses were repeated and 21-day-old females carrying both the mutation and the duplication were screened for motor axonal and NMJ phenotypes using confocal microscopy.

### Sanger sequencing confirmation

To confirm the presence of the disease-causing mutation in each mutant, sanger sequencing was performed on heterozygous mutant flies (FRT19A*/FRT19A). Phusion Plus DNA polymerase (F630L, Thermo Fisher) was used to amplify each fragment (see Table S3 for oligonucleotide sequences) before PCR cleanup (GeneJET Gel Extraction and DNA cleanup Micro Kit, Thermo Scientific) and sanger sequencing.

### Motor performance

To assess motor performance, a Rapid Iterative Negative Geotaxis (RING) assay was performed as described previously (Niehues *et al*, 2015). This assay is based on the innate escape response of flies to climb up the wall of a vial after being tapped down to its bottom. Flies are transferred into test tubes without anesthesia and five or six iterative measurements are video recorded on a Nikon Z50 camera.

#### *peb* RNAi and *Nf1* mutant motor performance

For analysis of *peb* RNAi and *Nf1^E1/+^* motor performance, *OK371-GAL4* virgins were collected and crossed to control, RNAi, or *Nf1^E1^* males. From those crosses, males were collected and housed with 6-10 flies per vial. Because flies stopped climbing around 21 days of age, speed was not an accurate measure. Therefore, we manually counted the number of flies per vial that crossed a 1.5 cm mark within 9 seconds versus flies that did not cross. Per vial a percentage of flies crossing was calculated for three videos, and averaged to obtain one data point for the respective vial. Per timepoint, at least 8 vials were analyzed per genotype.

#### Motor performance to characterize heterozygous *peb* mutants

For characterization of dominant motor performance deficits, mutant FRT19A*/FM7i virgins were crossed to FRT19A/Y males. FRT19A*/FRT19A virgins were collected in groups of 10 per vial. 10 to 15 vials of 1-day-old and 21-day-old flies were tested per genotype. Analysis was performed as described above.

### Lethal stage determination

For determination of the developmental stage of lethality, all mutant stocks were rebalanced with an *FM7c, twist-GAL4*>UAS-eGFP balancer (BL6873), that expresses eGFP in embryonic stages. 100-150 flies per genotype were kept O/N in a cage for egg-laying on an agar plate supplemented with yeast paste at 25 °C. The next day (day 2), ∼100 GFP-positive and ∼100 GFP-negative embryos were transferred to petri dishes containing Jazz-mix *Drosophila* medium (AS153, Fisher Scientific). On day 5, 2^nd^ instar larvae were counted and transferred to fresh medium, and on day 7 3^rd^ instar larvae were counted and transferred to regular food vials. Total number of pupae and eclosing adults were counted in the subsequent days. FRT19A was used as control. A developmental stage was marked lethal if <5% of animals survived the corresponding stage.

### Fluorescent in Situ Hybridization (FISH)

Hybridization chain reaction (HCR) RNA-FISH (Molecular Instruments) was used to detect peb or *rl* and *VGlut1* mRNA. MARCM crosses were set with *peb* mutants, and non-balancer females were collected and aged for 21 days at 25 °C. A previously described protocol (Choi *et al*, 2016) was adapted for *Drosophila* adult VNCs. Briefly, adult VNCs were dissected in PBS-T (PBS pH 7.2, 0.1% Tween 20 (P1370, Sigma Aldrich)), and fixed for 20 min in ice-cold 4% Formaldehyde (FA; 52549, Sigma-Aldrich) in PBS, followed by 20 min in ice-cold 4% FA in PBS-T. Three 10-minute washes with PBS-T were performed. Subsequently, VNCs were incubated for 3 min in 50% MeOH (M/4000/17, Fisher Chemical)/ 50% PBS-T, 3 min PBS-T, 3 min 100% MeOH, 3 min 50%MeOH / 50% PBS-T, and 3 min PBS-T. Another 20 min of fixing in 4% FA in PBS-T was performed before four 10-min washes in PBS-T. Prehybridization was performed by 14 minutes incubation in 4 μg/ml protK (740396, Macherey-Nagel). ProtK was washed away by rinsing twice with PBS-T, followed by two 5-minute PBS-T washes. A final 20 min 4% FA in PBS-T fixing step was performed before five 5-minute washes with PBS-T. VNCs were then incubated for 30 min in preheated probe hybridization buffer (Molecular Instruments) at 37 °C. Per sample, 1.6 pmol of each probe set (*peb* or *rl* and *VGlut1*) was added to 200μl of probe hybridization buffer and incubated O/N at 37 °C. The following day, four 15-minute washes with probe wash buffer were performed at 37 °C, followed by two 5-minute washes in SSC-T (5x sodium chloride sodium citrate (15557044, Invitrogen), 0.1% Tween-20) at RT. VNCs were then incubated with amplification buffer for 10 minutes. 6 pmol of hairpin H1 and 6pmol of hairpin H2 per sample were prepared separately by heating at 95 °C for 90s and cooled down for 30 minutes in the dark. Hairpins were added to 100 μl amplification buffer, and samples were incubated O/N in the dark. Hairpins were then removed by two 5-minute, two 30-minute and one 5-minute wash with SSCT. Samples were mounted in Vectashield (H1000-10, Vector Labs) and imaged on a Leica Sp8 Liachroic confocal microscope using a 63x objective. For quantification, the GFP channel was used to draw a circle around the plasma membrane in the plane where the nucleus diameter was the largest, and the mean intensity within this area was measured for each fluorescent channel for 4-10 cells per animal. *peb* or *rl* intensity was normalized to *VGlut1* intensity per cell. Then, the average normalized intensity of all cells per animal was used to obtain one datapoint. For *peb*, the normalized intensity of control animals without addition of the probe was averaged and used for background subtraction. Subsequently, data were normalized to FRT19A control.

### Immunohistochemistry

#### Larval NMJ

*OK371-GAL4* (for anti-Dlg1) or *OK371-GAL4*, UAS-mCD8::GFP; UAS-mCD8::GFP (all other larval NMJ immunostainings) virgins were crossed to control or *peb* RNAi males. Third instar larvae were dissected in HL3 buffer (70 mM NaCl, 5mM KCl, 1.5 mM CaCl_2_, 20mM MgCl_2_, 10 mM NaHCO_3_, 5 mM trehalose, 115 mM sucrose, and 5 mM Hepes 5 (pH 7.2)), and fixed in Bouin’s (HT10132, Sigma-Aldrich) for 3 min (for anti-Dlg1 and anti-Brp) or fixed in 4% FA (52549, Sigma-Aldrich) in PBS, pH 7.2, for 30 min (for anti-Syn and anti-Rop). Three 15 min washes were performed in PBT (PBS pH7.2, 0.2% Triton-X-100 (T8787, Sigma-Aldrich)), before blocking in 10% goat serum (16210064, Gibco) in PBT for 1 h. Incubation with primary antibody – either anti-Dlg1 (mouse monoclonal 4F3; DSHB; 1:200), anti-Brp (mouse monoclonal nc82; DSHB; 1:200), anti-Synapsin (mouse monoclonal 3C11; DSHB; 1:200) or anti-Rop (mouse monoclonal 4F8; DSHB; 1:100) – was performed in blocking solution O/N at 4 °C. The following day, three 15-minute washes in PBT were followed by a 3 h incubation with secondary antibody, either Alexa Fluor goat anti-mouse-488 (A-11001; Thermo Fisher; 1:500) or Alexa Fluor goat anti-mouse-568 (A11004; Thermo Fisher; 1:500). Finally, samples were washed 3x 15 min and mounted in Vectashield (H1000-10, Vector Labs). Images were taken of muscle 8 in abdominal segment 5 using a Leica SP8 Liachroic confocal microscope with a 20x Plan-Apochromat objective.

For calculation of NMJ length, maximum intensity projections of z-stacks comprising the entire NMJ were used to measure synapse length in ImageJ. For quantification of active zones, ImageJ plugin ‘DeadEasy Synapse’ (Sutcliffe *et al*, 2013) was adjusted to work with two-channel confocal images as input and then used to calculate the volume occupied by Bruchpilot voxels in cubic microns.

#### Adult ventral nerve cord

21-day-old adult flies from either MARCM or RNAi crosses were put on ice, briefly submerged in 100% ethanol (20821.296, VWR chemicals) before VNCs were dissected in PBT (PBS pH 7.2, 0.2% Triton-X-100 (T8787, Sigma-Aldrich)). Samples were fixed on ice for 30 min in 4% FA (52549, Sigma-Aldrich) in PBS, pH 7.2. Three 15-minute washes were performed in PBT before blocking (PBT, 2% w/v BSA (A9647, Sigma Aldrich), 5% goat serum (16210064, Gibco)) for 2 h, followed by incubation in primary antibodies in blocking solution for ∼64 h at 4 °C. Samples were then washed 6 times in blocking solution for 20 min followed by a 2 h incubation with secondary antibodies in blocking solution. After another six 20 min washes, VNCs were mounted in Vectashield (H1000-10, Vector Labs) and imaged on a Leica Sp8 Liachroic confocal microscope using a 63x objective.

Primary antibodies used were: anti-Peb (Mouse monoclonal 1G9; DHSB; 1:5), anti-tERK (rabbit polyclonal 9102S; Cell Signaling; 1:50), anti-pERK (mouse monoclonal M8159; Sigma; 1:200), anti-polyglutamylated alpha-tubulin (mouse monoclonal GT335; VWR; 1:500), anti-acetylated alpha-tubulin (mouse monoclonal 6-11B-1; Sigma; 1:800) and anti-Ref2P antibody (rabbit polyclonal ab178440; Abcam; 1:400).

Secondary antibodies used are Alexa Fluor goat anti-mouse-647 (A21235; Invitrogen; 1:500), Alexa Fluor goat anti-mouse-568 (A11004, Invitrogen; 1:500), Alexa Fluro goat anti-rabbit-568 (A11011; Invitrogen; 1:500) and Alexa Fluor goat anti-rabbit-647 (A-21245; Invitrogen; 1:500).

For Peb immunostaining, background was subtracted based on a control without addition of the primary antibody. If the GFP signal in any of the immunostainings was variable, normalization against GFP was performed before normalization against controls.

For quantification of Ref(2)P (P62), the number of Ref(2)P positive puncta was counted in 6 cells per animal by going through all planes of that cell. This number was averaged per animal to obtain one datapoint.

For quantification of fluorescence intensity for all other antibodies, the GFP channel was used to draw a circle around the plasma membrane in the plane where the nucleus diameter was the largest, and the mean intensity within this area was measured for each fluorescent channel for 4-10 cells per animal. Average intensity of all measured cells per animal was used to obtain one datapoint.

#### TUNEL

For the TUNEL (Terminal deoxynucleotidyl transferase-mediated dUTP Nick End Labeling) assay, 21-day-old adults from MARCM crosses were put on ice, briefly submerged in ethanol before VNCs were dissected in PBT (PBS pH 7.2, 0.2% Triton X-100 (T8787, Sigma-Aldrich)). The protocol was adapted from Denton et al (Denton & Kumar, 2015). Briefly, samples were fixed at RT for 20 min in 4% FA (252549, Sigma-Aldrich) in PBS, pH 7.2. Three 10-minute washes were performed in PBT followed by one wash in 0.5PBT (PBS pH 7.2, 0.5% Triton-X-100). Permeabilization was performed in 100 mM sodium citrate (S279, Fisher Scientific, freshly prepared, in PBT). Three 5-minute washes in PBT were performed before preparing the TUNEL (In Situ Cell Death Detection Kit, TMR red, Roche) reaction according to manufacturer’s instructions. Briefly, we used 10 μl of enzyme and 90 μl of labelling mix per sample, and incubated for 3 h at 37°C. Finally, three 10-minute washes were performed in PBT before mounting VNCs in Vectashield (H1000-10, Vector Labs). VNCs were imaged on a Leica Sp8 Liachroic confocal microscope using a 40x objective. Control flies which had received a 1.5 h heat shock at 37°C were included as a positive control for TUNEL staining. For quantification, the total number of homozygous mutant clones per animal and the number of TUNEL positive homozygous mutant clones were counted. The percentage of positive cells was calculated per animal to obtain one data point.

### Autophagy flux sensor

Control and *peb* RNAi virgins were crossed to *OK371-GAL4*, UAS-GFP-mCherry-Atg8a/CyO males. 21-day-old non-balancer males were anesthetized on ice and briefly submerged in ethanol before VNCs were dissected in PBT (PBS pH 7.2, 0.2% Triton-X-100 (T8787, Sigma-Aldrich)). Samples were fixed on ice for 30 min in 4% FA (252549, Sigma-Aldrich) in PBS, pH 7.2. Three 10-minute washes were performed in PBS before mounting VNCs in Vectashield (H1000-10, Vector Labs). VNCs were imaged on a Leica Sp8 Liachroic confocal microscope using a 63x objective. Total number of GFP + mCherry (yellow, autophagosomes) and mCherry only (autolysosomes) puncta were counted in all planes for 6 cells per animal. The total number of Atg8a puncta was determined per cell and averaged per animal to obtain one data point. For calculation of autophagic flux, the total number of autolysosomes was divided by total number of autophagosomes per animal to obtain one data point.

For ex vivo treatment with Bafilomycin A1, VNCs were dissected as described above. Dissected VNCs were kept for 6 h at 25 °C in either HL3 buffer (70 mM NaCl, 5mM KCl, 1.5 mM CaCl_2_, 20mM MgCl_2_, 10 mM NaHCO_3_, 5 mM trehalose, 115 mM sucrose, and 5 mM Hepes 5 (pH 7.2)), or HL3 buffer containing 200 nM Bafilomycin A1 (HY-100558, MedChemExpress). After treatment, samples were fixed on ice for 30 min in 4% FA (252549, Sigma-Aldrich) in PBS, pH 7.2. Three 10-minute washes were performed in PBS before mounting VNCs in Vectashield (H1000-10, Vector Labs). VNCs were imaged on a Leica Sp8 Liachroic confocal microscope using a 63x objective. Quantification was performed as described above.

### Mushroom body MARCM

Control (FRT19A) or *peb* mutant virgins were crossed to hs-FLP, tubP-GAL80, FRT19A; UAS-mCD8::GFP/CyO; *OK107-GAL4* males. Heat shock was performed for 1h at 37 °C 6 days after hatching. Female offspring was collected and aged for 21 days before brains were dissected in PBT (PBS, 0.2% Triton-X-100, pH7.2) and subsequentially fixed for 30 min in ice-cold 4% FA (252549, Sigma-Aldrich) in PBS, pH 7.2.. Three 10-minute washes were performed in PBT, before blocking for 1h in 10% goat serum in PBT on a rocking platform. Samples were incubated with rabbit anti-GFP (Invitrogen; A6455; 1:200) in blocking buffer O/N at 4°C. Another three 10-minute washes were performed in PBT before incubation for 3h in goat anti-rabbit-488 secondary antibody (Invitrogen; A11008; 1:500). Finally, three 10-minute washes in PBS were performed, before mounting in Vectashield (H1000-10, Vector Labs). Mushroom body β-lobes were imaged on a Leica Sp8 Liachroic confocal microscope using a 63x objective.

### Drug feeding experiments

*OK371-GAL4* virgins were crossed to control or *peb* RNAi males to obtain offspring for motor performance assays, while mutant virgins were crossed to FRT19A, Tub-GAL80*; OK371-GAL4*, UAS-mCD8::GFP; ase-FLP, UAS-mCD8::GFP males. Newly eclosed males (for RNAi crosses) or females (for MARCM crosses) were collected and 1-day-old flies were transferred to regular food or Jazz mix food (AS153, Fisher Scientific) prepared according to manufacturer’s instructions, containing either the drug-of-interest or an identical quantity of solvent (vehicle). Flies were weekly transferred to freshly prepared food.

For Mirdametinib treatment, mirdametinib (HY10254G, MedChemExpress) was dissolved in 100% ethanol (20821.296, VWR chemicals) and used at a final concentration of 0.1 or 1 μM. For Tubastatin A (HY-13271A, MedChemExpress) treatment, Tubastatin A was dissolved in DMSO (D2438, Sigma-Aldrich), and used in a final concentration of 10 μM. For Rapamycin (HY-10219, MedChemExpress) treatment, DMSO was used as a solvent and a final concentration of 50 μM was used.

### Temporal peb RNAi expression

For expression of *peb* RNAi selectively during development, adulthood or both, *OK371-GAL4*, UAS-mCD8::GFP, tub-GAL80^ts^ virgins were crossed to *peb* RNAi 1 or control males. For developmental expression of the RNAi, crosses were set at 29°C. Newly eclosed male flies were transferred to 18°C to switch off UAS transgene expression. For selective expression in adult flies, crosses were set at 18°C and newly eclosed male flies were transferred to 29°C. For continuous expression, crosses were set at 29°C and newly eclosed male flies were kept at 29°C. For developmental expression of the RNAi, 19-day-old males were transferred to 29°C to allow for UAS-mCD8::GFP expression. Legs of 21-day-old males were dissected, mounted in Prolong Glass antifade mountant (P36980, Thermo Scientific) and imaged on a Leica Sp8 Liachroic confocal microscope using a 63x objective.

### Quantification and statistical analysis

All results of statistical analysis are presented as mean ± standard error of the mean. Differences were considered significant when p<0.05. No statistical methods were used to pre-determine sample sizes. Whenever possible, data collection and analysis were performed by investigators blinded to the genotype of the experimental animals.

Before analysis, a Robust regression and Outlier removal method was performed to detect and exclude outliers. Assumptions of normality were tested by Shapiro-Wilk, Anderson-Darling and Kolmogorov-Smirnov tests, whereas assumptions on homogeneous variances (heteroscedasticity) were tested by Brown-Forsythe and Barlett’s test. Statistical tests were only performed if all their assumptions were met, the statistical test used per panel is described in each figure legend. Statistical analysis was performed using GraphPad Prism v10.2.1.

For comparison of normally distributed data between two groups, a two-tailed unpaired t-test was performed. When normality assumptions were not met, a Mann-Whitney test was performed.

For comparisons of normally distributed data between more than two groups and one factor (genotype), One-Way ANOVA with Tukey test for multiple comparisons was performed. If normality assumptions were not met, Kruskal-Wallis with Dunn’s multiple comparisons test was used. If heteroscedasticity assumptions were not met, a Brown-Forsythe and Welch ANOVA with Dunnett’s T3 multiple comparisons test was used.

For comparisons between more than two groups with two factors (genotype and treatment), two-way ANOVA with Geisser-Greenhouse correction and Tukey multiple comparison test was used. If normality assumptions were not met, Kruskal-Wallis with Dunn’s multiple comparisons test was used. If heteroscedasticity assumptions were not met, a Brown-Forsythe and Welch ANOVA with Dunnett’s T3 multiple comparisons test was used. For longitudinal motor performance data, normality and heteroscedasticity assumptions were tested individually for each relevant comparison.

## Acknowledgements

We thank Marc Freeman, Lukas Neukomm and Mikio Furuse for providing *Drosophila* lines, the General Instruments Facility (Faculty of Science, Radboud University) for advice on image acquisition and analysis and Marijn Kuijpers for advice on Atg8a flux sensor analysis. We thank Youp van Oosterhout, Anna Bolhuis, Hanna Vreemann, Andrea Mlinar, Bram van Raalte, Yujin Hong, Vasilis Kougioumzoglou, Paulien Sloot, Nikki Smolders, Jimmy Vu, Kiki Peeters, Gino Hulshof, Dennis Colin and Céline van Diermen for their help during their internships. M.B. and E.S. were supported by a grant from ‘Stichting ALS Nederland’, E.S. received an ERC consolidator grant (ERC-2017-COG 770244), and funding from the Radala Foundation, the ‘Prinses Beatrix Spierfonds’ (W.OR22-03), the Muscular Dystrophy Association (MDA 946876), AFM-Telethon, ARSLA, an NWO Open Competition ENW-M grant, an NWO Take-off WO Phase 1 grant, and a grant from the DI-NIN Fund.

## Author contributions

Conceptualization, M.B. and E.S.; Methodology, M.B and E.S.; Software, M.B.; Validation, M.B.; Formal analysis, M.B, S.H., P.P., D.B., G.A. and K.K.; Investigation, M.B., E.L., A.G., A.Se., A.B., S.H., M.K., C.M., P.P., H.R.A., B.R., S.G., P.L. and C.S.; Resources, P.V, J.V, S.Z., C.G., S.L.A., A.Sc. and E.S.; Data curation, M.B.; Writing – Original Draft, M.B.; Writing – Review & Editing, M.B., D.B., B.D., P.V., J.V., S.Z., C.G., S.L.A., A.Sc. and E.S.; Visualization, M.B. and E.S.; Supervision, A.Sc. and E.S.; Funding Acquisition, M.B. and E.S.

## Data availability

Raw data obtained in this study have been deposited in the Radboud repository.

---

**Figure S1:**
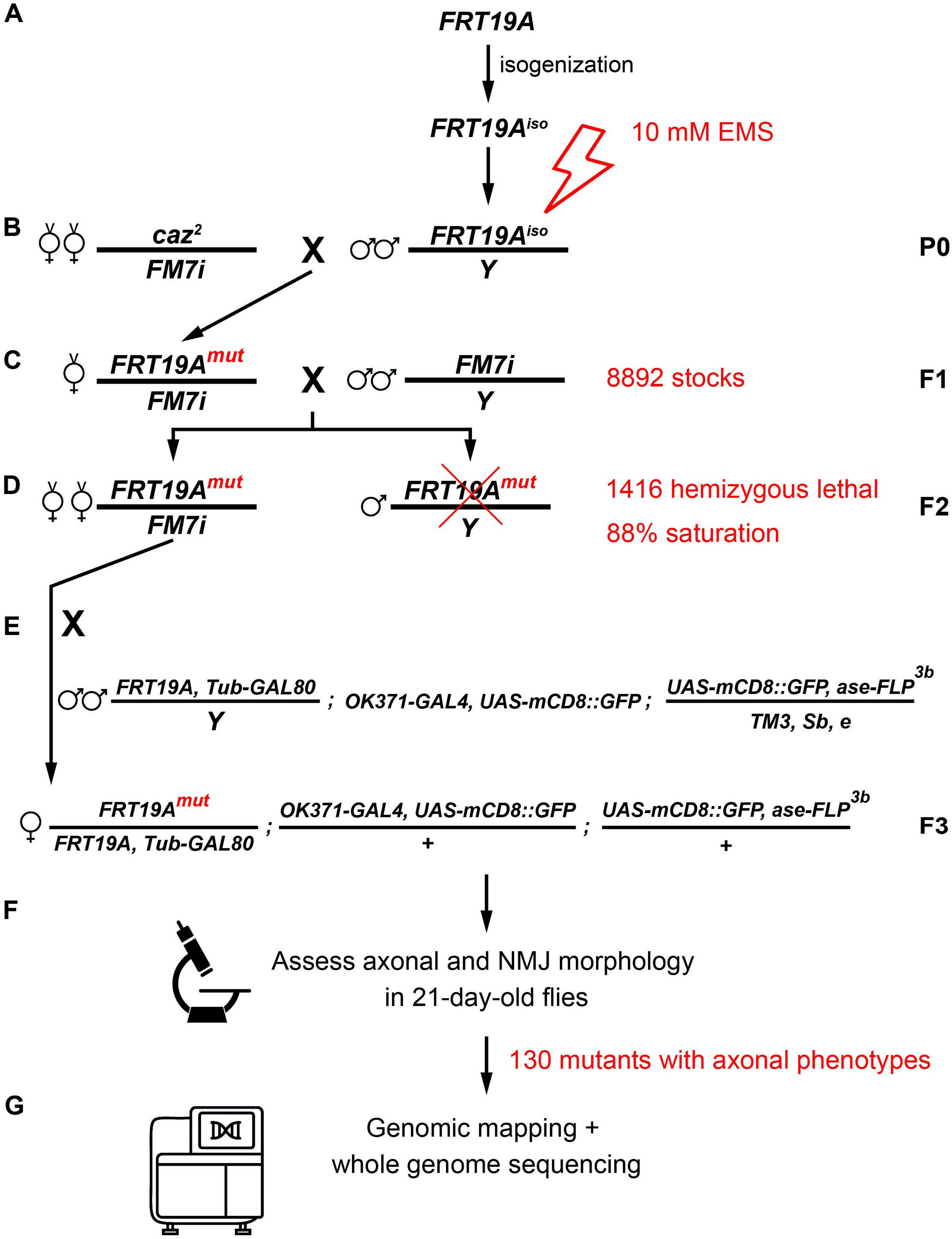
Schematic overview of the F3 adult mosaic genetic screen of the *Drosophila* X chromosome. **(A)** The FRT19A chromosome was isogenized. Male flies were mutagenized and **(B)** crossed to balancer virgins to obtain **(C)** FM7i balanced females with unique mutations. **(D)** Stable stocks were built and assessed for hemizygous lethality. 1416 X-linked recessive lethal stocks were identified and **(E)** crossed to MARCM males to **(F)** assess axonal and NMJ morphology in 21-day-old flies. **(G)** 130 mutants with motor axonal phenotypes were identified (including putative cell-lethal mutants in which no GFP-labeled axons or NMJs could be observed) and used for genomic mapping and whole genome sequencing to identify the causative gene. *caz^2^/FM7i* virgins were used because the *caz^2^* allele is homozygous lethal (Frickenhaus *et al.*, 2015).

**Figure S2:**
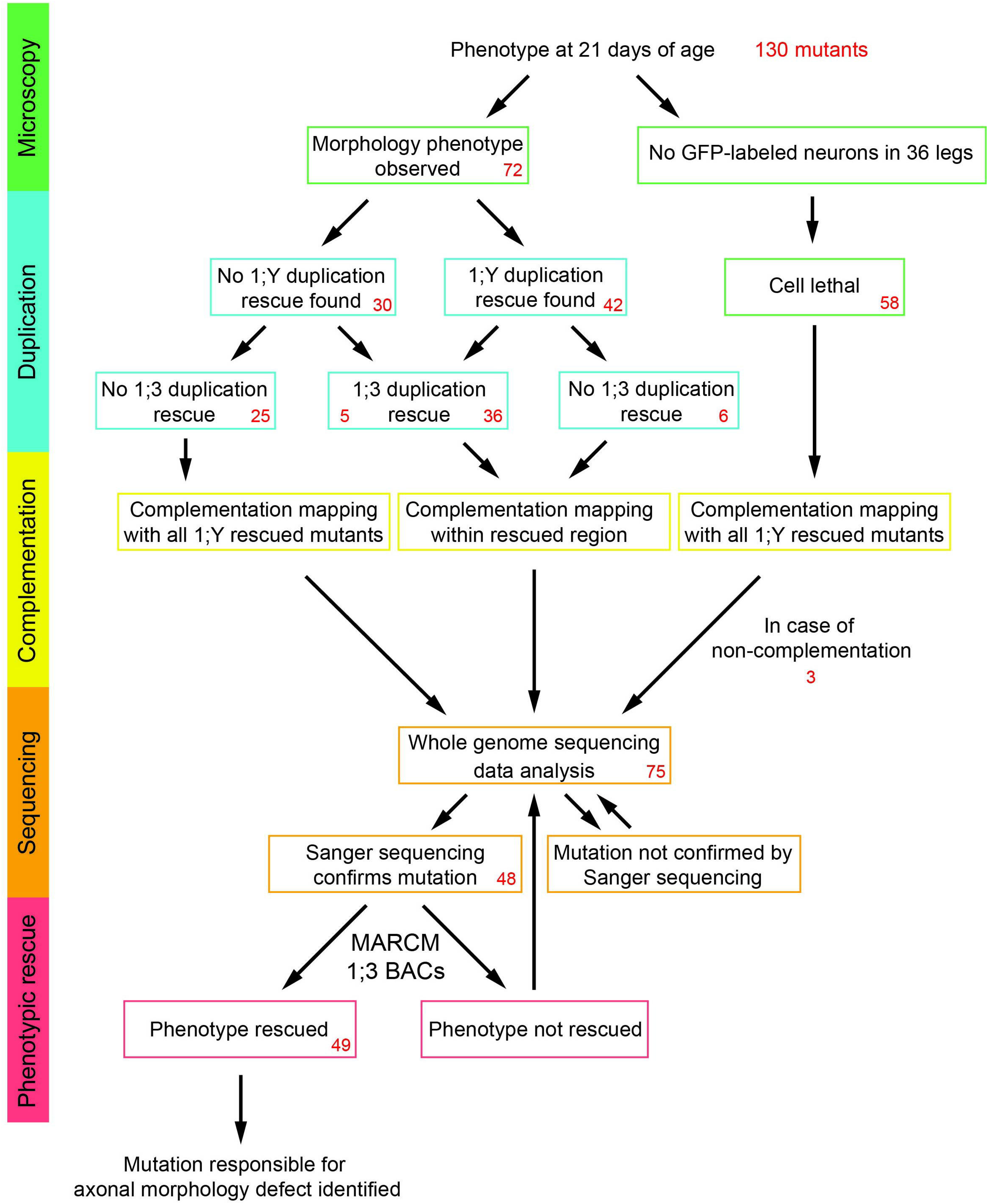
Flow chart showing the steps towards identification of the causative mutations in identified neurodegeneration mutants. The subsequent steps involving microscopy analysis, duplication mapping, complementation mapping, whole genome and Sanger sequencing, and phenotypic rescue experiments are shown. The number of mutants in each step is indicated in red.

**Figure S3:**
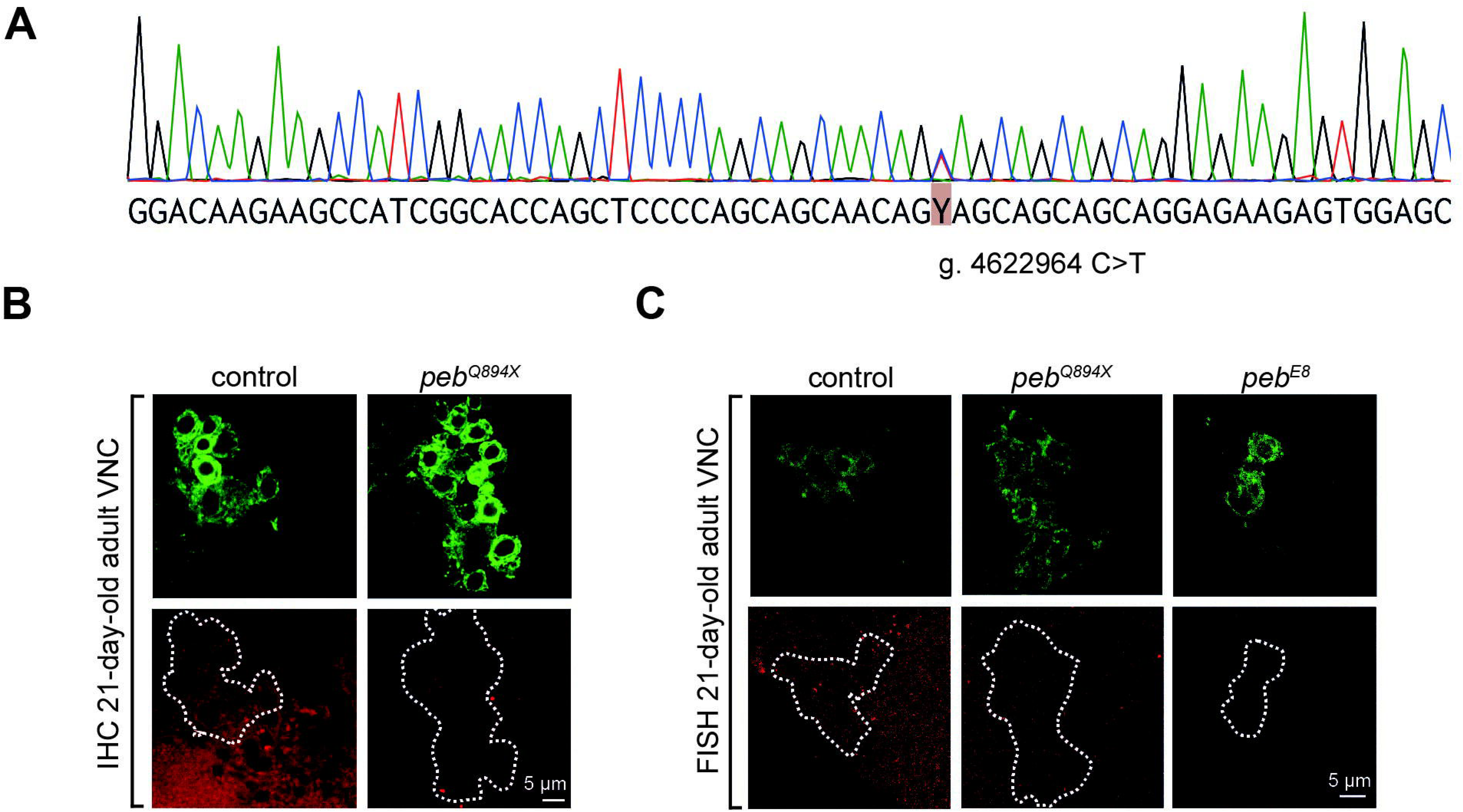
Characterization of *peb* loss-of-function alleles. **(A)** Sanger sequencing results to confirm the *peb^Q894X^* heterozygous mutation. **(B,C)** Representative images of (B) Peb immunohistochemistry and (C) FISH for peb transcript in motor neuron cell bodies in the VNC of 21-day-old adult flies. *OK371-GAL4*-driven GFP expression labels MARCM-induced homozygous mutant motor neuron clones. Scale bars: 5 μm.

**Figure S4:**
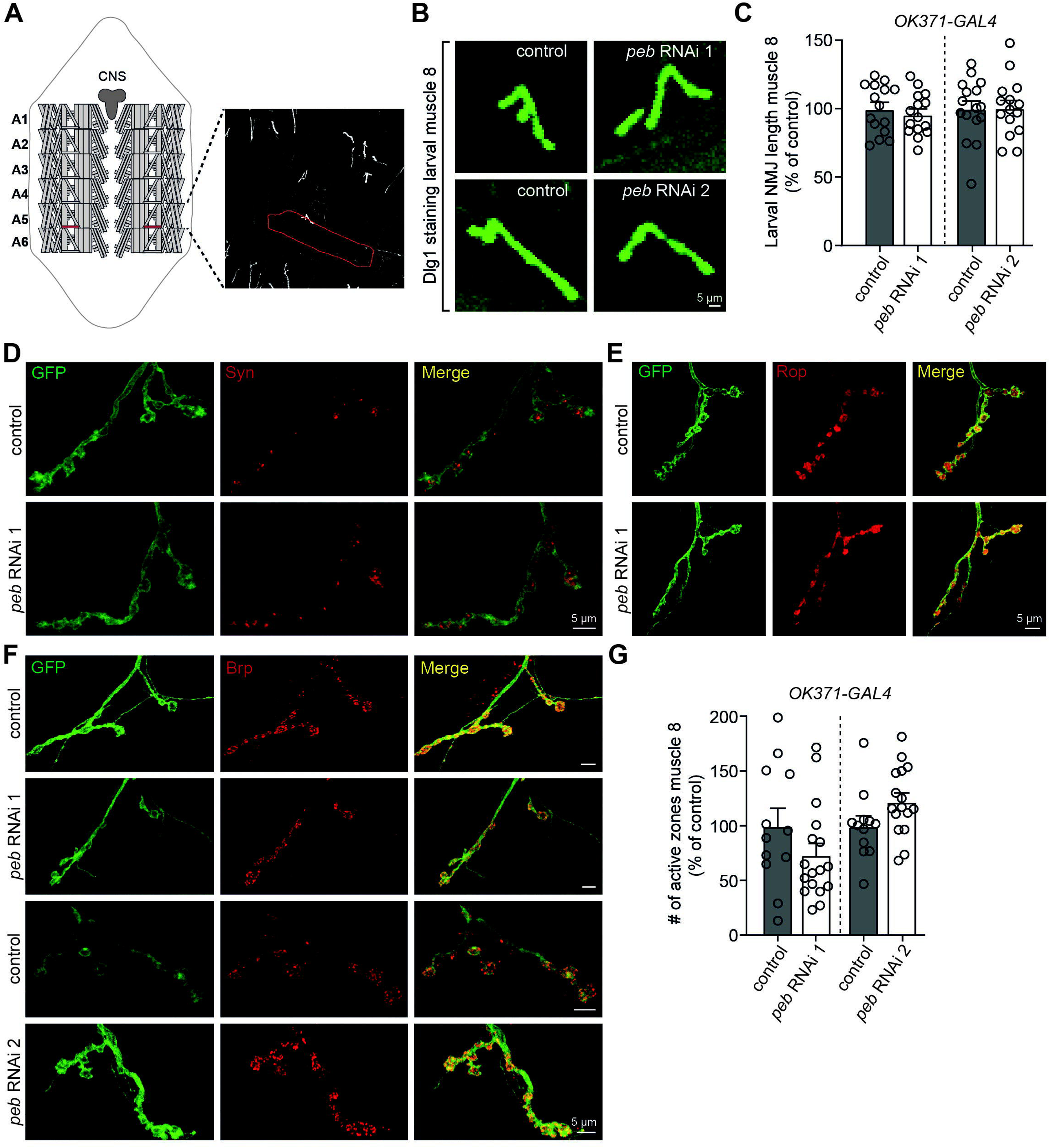
Loss of *peb* function results in an adult-onset phenotype. **(A)** Overview of the muscle architecture in the larval body wall. Imaging and quantification was performed in muscle 8 in abdominal segment 5 of third instar larvae (delineated in red). **(B)** Representative images and **(C)** quantification of NMJ length in larvae with *peb* knockdown in motor neurons (*OK371-GAL4*) using two independent RNAi lines versus genetic background controls (KK or GD background, respectively). NMJs were visualized using immunostaining for the postsynaptic marker Dlg1. **(D-F)** Representative images of immunostaining for presynaptic markers, including (D) synaptic vesicle protein Synapsin (Syn), (E) SNARE-binding protein Ras opposite (Rop) and (F) active zone marker Bruchpilot (Brp) on the NMJ on larval muscle 8, in the presence or absence of *peb* knockdown in motor neurons (*OK371-GAL4*). **(G)** Quantification of the number of active zones in the larval NMJ after *peb* knockdown in motor neurons (*OK371-GAL4*) using two independent RNAi lines versus genetic background controls (KK or GD background, respectively). P=not significant by (C,G) unpaired t-test. n= (C) 15-16 animals per genotype and (G) 12-17 animals per genotype. Data are represented as mean ± SEM. Scale bars: 5 μm.

**Figure S5:**
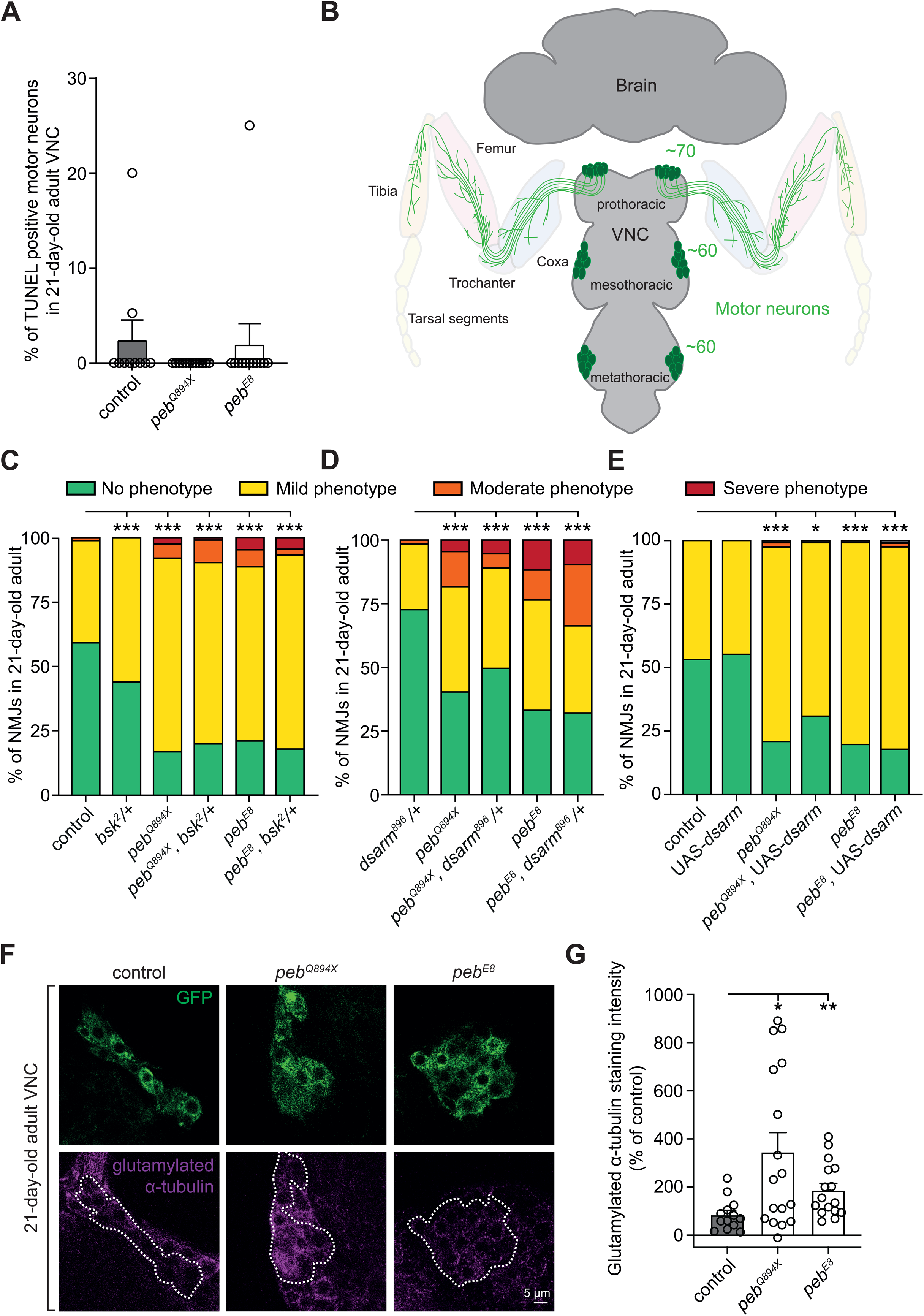
*peb* mutant phenotypes are independent of Wallerian degeneration, cell death pathways or reductions in glutamylated α-tubulin. **(A)** Quantification of the percentage of TUNEL positive nuclei in *peb* mutant versus control (FRT19A) motor neuron clones in 21-day-old adult VNCs. **(B)** Schematic overview of motor neuron clusters in the *Drosophila* ventral nerve cord (VNC). Each leg is innervated by 60-70 motor neurons(Guan *et al.*, 2018). **(C)** Semi-quantitative analysis of femoral NMJ degeneration in control (FRT19A) and *peb* mutant motor neuron clones of 21-day-old animals in the presence or absence of a heterozygous loss-of-function mutation in the JNK ortholog *bsk.* (D,E) Semi-quantitative analysis of femoral NMJ degeneration in control and *peb* mutant motor neuron clones in 21-day-old adult flies, in the presence or absence of (D) a heterozygous *dsarm* mutation, or (E) UAS-*dsarm*. **(F)** Representative images and **(G)** quantification of immunostaining intensity of glutamylated α-tubulin in cell bodies of *peb* mutant versus control (FRT19A) motor neuron clones in 21-day-old adult VNCs. Data are shown as percentage of control. Scale bar in (F): 5 μm. *P<0.05, **P<0.01, ***P<0.005 by (A) Kruskal-Wallis test with Dunn’s multiple comparisons test, (C,D,E) Fisher’s exact test with Bonferroni correction or (G) Brown-Forsythe and Welch ANOVA with Dunnett’s T3 multiple comparisons test. n= (A) 10-12 animals per genotype, (C) 73-114 NMJs in 6-10 animals per genotype, (D) 38-141 NMJs in 6-10 animals per genotype, (E) 86-108 NMJs in 6-10 animals per genotype and (G) 13-17 animals per genotype. Data are represented as mean ± SEM.

**Figure S6:**
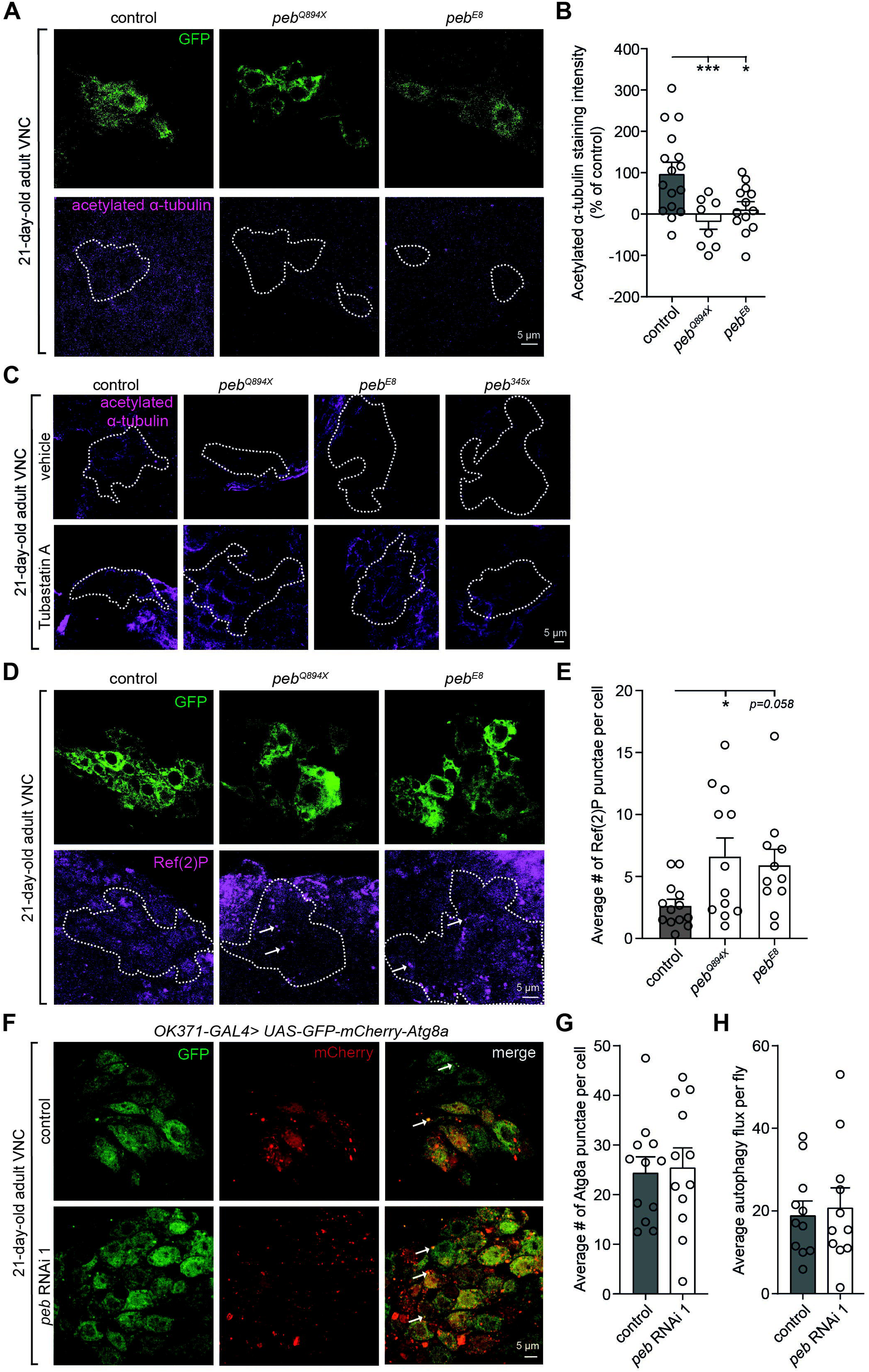
Loss of *peb* function results in defects in α-tubulin acetylation and autophagy. **(A)** Representative images and **(B)** quantification of immunostaining intensity of acetylated α-tubulin in control (FRT19A) versus *peb* mutant motor neuron clones in the VNC of 21-day-old flies. Data was normalized to control without primary antibody, and shown as percentage of control (FRT19A). **(C)** Representative images of immunostaining intensity of acetylated α-tubulin in control (FRT19A) versus *peb* mutant motor neuron clones in the VNC of 21-day-old flies treated with 10 μM Tubastatin A or vehicle. **(D)** Representative images and **(E)** quantification of Ref(2)P-positive punctae (indicated by arrows in D) in *peb* mutant versus control (FRT19A) motor neuron clones in the VNC of 21-day-old flies. **(F)** Representative images and **(G)** quantification of the average number of mCherry-positive Atg8a puncta per motor neuron (*OK371-GAL4*) in the ventral nerve cord of 21-day old flies expressing the GFP-mCherry-Atg8a autophagy marker, with or without simultaneous *peb* knockdown. **(H)** Quantification of autophagic flux, calculated by dividing the total number of red puncta (autolysosomes) per animal by the total number of yellow puncta (autophagosomes) per animal. *P<0.05, ***P<0.005 by (B) one-way ANOVA with Dunnett’s multiple comparisons test or (E) Brown-Forsythe and Welch ANOVA with Dunnett’s T3 multiple comparisons test or (G,H) two-tailed unpaired t-test. n= (B) 8-16 animals per genotype, (E) 11-13 animals per genotype and (G,H) 11-12 animals per genotype. Data are represented as mean ± SEM. Scale bars: 5 μm.

**Figure S7:**
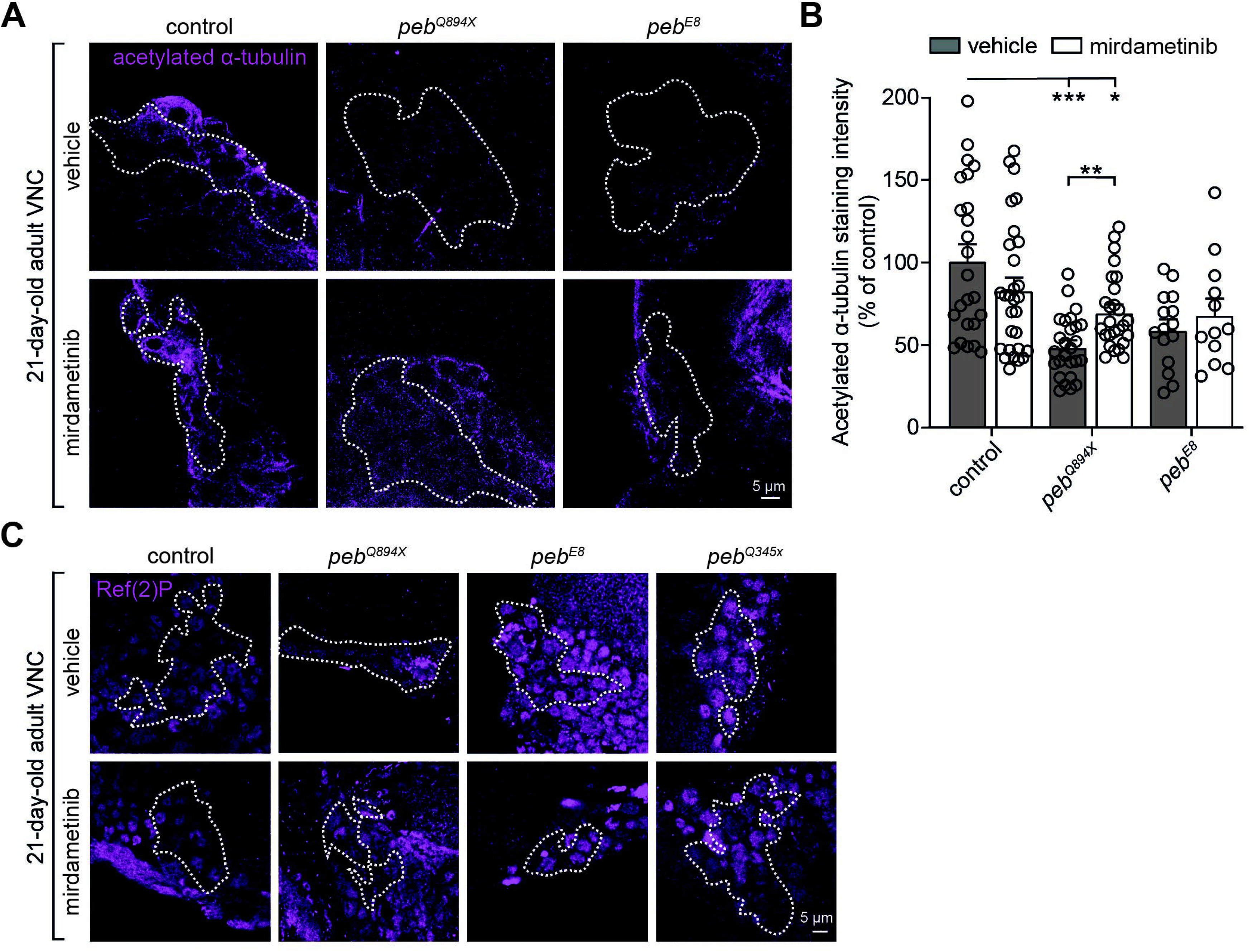
Defects in α-tubulin acetylation and autophagy are downstream of excessive RAS/MAPK signaling. **(A)** Representative images and **(B)** quantification of immunostaining intensity of acetylated α-tubulin in control (FRT19A) versus *peb* mutant motor neuron clones in the VNC in 21-day-old flies treated with 1μM mirdametinib or vehicle. Data were normalized to control. **(C)** Representative images of Ref(2)P-positive puncta in *peb* mutant versus control (FRT19A) motor neuron clones in the VNC of 21-day-old flies, treated with 1μM mirdametinib or vehicle. *P<0.05, **P<0.01, ***P<0.005 by (B) Brown-Forsythe and Welch ANOVA with Dunnett’s T3 multiple comparisons test. n= (B) 12-29 animals per experimental group. Data are represented as mean ± SEM. Scale bars: 5 μm.

